# Hippocampal Ripple Diversity organises Neuronal Reactivation Dynamics in the Offline Brain

**DOI:** 10.1101/2025.03.11.642571

**Authors:** Manfredi Castelli, Vítor Lopes-dos-Santos, Giuseppe P. Gava, Renaud Lambiotte, David Dupret

## Abstract

Hippocampal ripples are highly synchronized neuronal population patterns reactivating past waking experiences in the offline brain. Whether the level, structure, and content of ripple-nested activity are consistent across consecutive events or are tuned in each event remains unclear. By profiling individual ripples using laminar currents in the mouse hippocampus during sleep/rest, we identified Rad^sink^ and LM^sink^ ripples featuring current sinks in *stratum radiatum* versus *stratum lacunosum-moleculare*, respectively. These two ripple profiles recruit neurons differently. Rad^sink^ ripples integrate recent motifs of waking coactivity, combining superficial and deep CA1 principal cells into denser, higher-dimensional patterns that undergo hour-long stable reactivation. In contrast, LM^sink^ ripples contain core motifs of prior coactivity, engaging deep cells into sparser, lower-dimensional patterns that undergo a reactivation drift, gradually updating their pre-structured content for recent wakefulness. We propose that ripple-by-ripple diversity instantiates parallel reactivation channels for stable integration of recent wakefulness and flexible updating of prior internal representations.

## Introduction

The brain has a remarkable ability to retain prior knowledge while continuously integrating new information. Understanding how neuronal populations coordinate these two operations, enabling the seamless updating of brain networks, remains a fundamental biological question. The offline states of sleep and rest could play a key role in this process by providing time windows for the reorganization and stabilization of neuronal activity patterns ^1–3^. Notably, hippocampal ripples have emerged as critical brain network events for facilitating offline processing of neuronal population activity ^4,5^.

Hippocampal ripples are transient network events in the mammalian brain, characterized by high-frequency oscillations (100–250 Hz) in the local field potentials (LFPs) of the CA1 pyramidal cell layer (*stratum pyramidale*). These events, prevalent during sleep and rest, are paired with a prominent negative deflection in the LFPs of the *stratum radiatum* – the sharp wave – reflecting inputs from CA3 to CA1 ^4,6^. The sharp-wave/ripple (SWR) complex has a clear electrophysiological signature, with a well-defined laminar current source density (CSD) profile featuring a sink in the CA1 *stratum radiatum* ^4^. Hippocampal ripples represent the most synchronous pattern of neuronal population activity in the mammalian brain, providing discrete time windows during which hippocampal principal cells discharge action potentials collectively. This synchrony facilitates the “reactivation” of coactivity patterns from waking experiences for further offline processing during sleep and rest periods ^7–15^.

The functional relevance of ripples has been demonstrated in studies showing that their suppression destabilizes hippocampal activity patterns and impairs memory performance ^16–20^, while promoting ripples and their associated spiking activity improves memory retention ^21,22^. Beyond their canonical role in memory, ripple-nested spiking is also linked to other behavioural functions, including planning, inference, and even metabolism ^9,10,23–25^. Moreover, ripples provide a framework for brain-wide interactions, with both intra- and extra-hippocampal inputs shaping their electrophysiological features ^26,27^. Yet, since their discovery, hippocampal ripples have typically been treated as a homogenous network pattern, raising the question of how ripple-by-ripple variability could possibly support diverse functions. Recent evidence suggests that hippocampal ripples are not homogeneous events but exhibit significant variability in their electrophysiological expression, forming a continuum of features in the LFP waveform space ^28,29^. This variability may arise from differences in how neural inputs are organised along the somato-dendritic axis of CA1 principal cells ^30,31^. However, this diversity remains underexplored because traditional ripple analyses rely on averaging spectral characteristics, potentially masking important differences in these network events ^27–29^. Identifying the ripple-to-ripple variability in activity levels, structural organization, and neuronal content of population patterns is essential for understanding how these brief events support hippocampal computations.

To investigate hippocampal ripple diversity, we explored the current source density underlying each individual ripple across the CA1 neural layers. This revealed two laminar profiles that we refer to as Rad^sink^ versus LM^sink^ ripples, for *stratum pyramidale* events paired with a current sink in *stratum radiatum* versus those with a current sink in *stratum lacunosum-moleculare*. These two profiles engage CA1 and CA3 neurons differently, host distinct motifs of millisecond-timescale coactivity, and exhibit distinct topological organisation of their population patterns. LM^sink^ ripples contain small motifs of stronger coactivity between principal cells in the deep CA1 pyramidal sublayer. Following waking experience, Rad^sink^ ripples append onto these core motifs the recently recruited superficial CA1 principal cells, yielding denser, higher-dimensional population patterns that consistently reactivate over hour-long sleep/rest. Meanwhile, the pre-existing coactivity backbone nested in LM^sink^ ripples, formed by sparser, lower-dimensional patterns, gradually drifted throughout sleep/rest, slowly updating its content to reflect recent wakefulness. Collectively, these findings reveal two ripple profiles with distinct population activity levels, structural organization, and neuronal content that cooperate to balance the integration of recent experiences with the updating of neuronal priors in the offline hippocampus.

## Results

### Profiling individual hippocampal ripples using their laminar currents

We started to investigate the diversity of hippocampal ripples by simultaneously recording LFPs in each layer along the somato-dendritic axis of CA1, from *stratum oriens* to the hippocampal fissure, using silicon probes (Table S1). This recording setup enabled both ripple detection in the LFPs of *stratum pyramidale* and Current Source Density (CSD) analysis of laminar currents during sleep/rest (Figures 1A-C and S1A; 52.2 total hours of sleep/rest from 5 mice; 38 sleep/rest sessions; mean duration (IQR): 82.4 (59.5 – 105.9) minutes per sleep/rest session). In line with existing knowledge ^4^, the average ripple profile displayed a current sink in *stratum radiatum* and a current source in *stratum lacunosum-moleculare* (Figure 1A). However, individual ripples showed considerable variability in their CSD profiles (Figures 1B and S1B,C). Most ripples featured a current sink in *stratum radiatum* (Figures 1B and S1B), reflecting the grand average CSD profile (Figure 1A). The strength of this *stratum radiatum* sink positively correlated with the strength of a concurrent sink in *stratum oriens* (Figure S1D,E). In addition to these canonical events, other ripples exhibited a strong current sink in *stratum lacunosum-moleculare*, accompanied by a source in the *stratum radiatum* (Figures 1B and S1C-E), diverging from the grand average CSD profile (Figure 1A).

**Figure 1:**
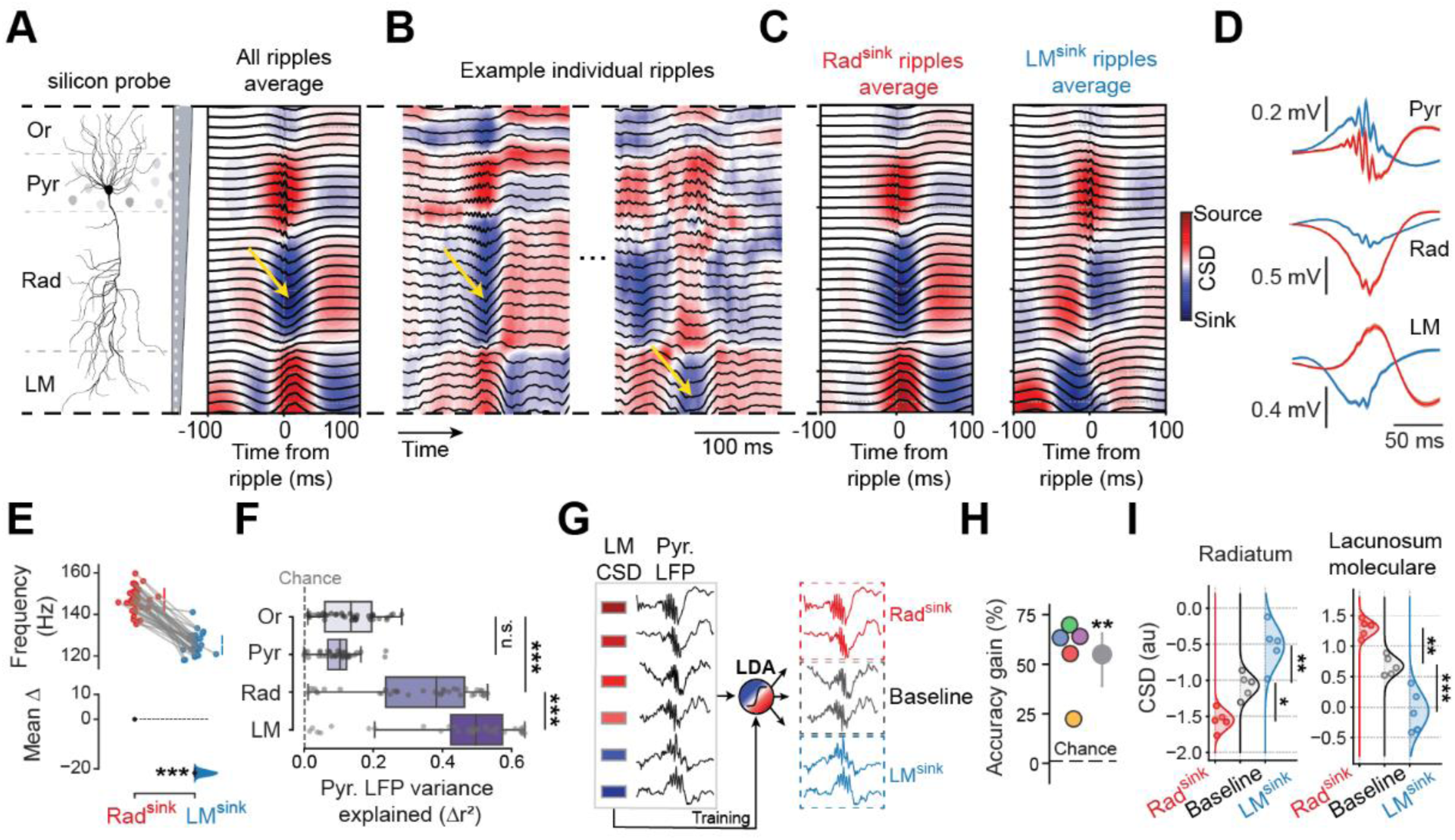
Laminar current profiles reveal two types of hippocampal CA1 ripples. **(A)** Average current source density (CSD; colour-coded map) and local field potential (LFP) waveform (black traces) for ripples recorded in *stratum pyramidale* using a silicon probe spanning the parallel neural layers of CA1 (Or, *oriens*; Pyr, *pyramidale*; Rad, *radiatum*; LM, *lacunosum-moleculare*). *Yellow arrow*, current sink in *stratum radiatum*. **(B)** Instantaneous CSD and LFP waveform for two example ripples from one sleep session. **(C)** Average CSD and LFP waveform for ripples with a strong current sink either in *radiatum* (Rad^sink^ ripples) or *lacunosum-moleculare* (LM^sink^ ripples). **(D)** Average waveform for Rad^sink^ (red) and LM^sink^ (blue) ripples using LFPs simultaneously recorded in the pyramidal (Pyr), radiatum (Rad) and lacunosum-moleculare (LM) layers. To visualise differences in ripple frequencies, LFP waveforms were referenced to the highest ripple-peak. *Shaded area* around each trace, 95% CI. **(E)** Estimation plot showing the effect size for the difference in average ripple frequency between Rad^sink^ and LM^sink^ ripples. *Top panel*, raw data distributions (each dot represents a sleep session). *Bottom panel*, mean difference (*black dot*, mean; *black ticks*, 95% confidence intervals; *coloured area*, bootstrapped error distribution). **(F)** Variance explained in the ripple LFP signal of the pyramidal layer by the CSD in each CA1 layer. Each dot represents a sleep session. Note that the lacunosum-moleculare CSD explains most of the variance (p = 1.2×10^-24^, one-way ANOVA; Rad versus LM: p = 5.7×10^-4^, Tukey post-hoc). **(G)** Schematic of linear discriminant analysis (LDA) predicting the CSD in *lacunosum-moleculare* from the ripple LFP signals in *stratum pyramidale*. Each ripple LFP trace and corresponding sign and magnitude of its CSD signal used in the LDA classifier to distinguish Rad^sink^, baseline and LM^sink^ ripples. **(H)** LDA accuracy. Each dot represents one mouse left outside the training set (leave-one-out approach). *Dashed line*, chance level; *gray dot*, mean; *vertical ticks*, 95% CI. **(I)** Radiatum and lacunosum-moleculare CSD values in LDA-classified Rad^sink^, baseline, and LM^sink^ ripples. For each leave-one-out model, we applied the trained model to the mouse left-out of training and computed the mean CSD in each predicted ripple. Each dot represents one mouse left-out of training. LDA models significantly discriminated ripples with a strong radiatum current sink, those with a baseline CSD profile, and those with a strong lacunosum-moleculare sink (radiatum CSD, p = 2.6×10^-5^; lacunosum-moleculare CSD, p = 3.1×10^-6^; one-way ANOVA with Tukey post-hoc) *p < 0.05, **p < 0.01, ***p<0.001.

We studied ripple variability using each event’s CSD signature, defined as the CSD signal within a 50-ms window centred on its ripple power peak. This approach allowed representing each ripple as a curve characterizing its laminar current profile along the CA1 somato-dendritic axis (Figure S1B,C). Applying principal component analysis to the distribution of individual CSD signatures unveiled two distinct laminar profiles that delineate a continuum (Figures 1C and S2A-E). The first profile showed a stronger current sink in *stratum radiatum* (Rad); we hereafter refer to these events as Rad^sink^ ripples (Figure 1C). The second profile showed a stronger current sink in *stratum lacunosum-moleculare* (LM); we hereafter refer to these events as LM^sink^ ripples (Figure 1C). In terms of LFP waveforms, Rad^sink^ ripples exhibited the well-described negative *radiatum* sharp-wave, while LM^sink^ ripples had a pyramidal layer positive deflection with a *lacunosum-moleculare* negative deflection (Figure 1D). Compared to Rad^sink^, LM^sink^ events exhibited lower ripple frequency[Figures 1E and S2F-H; mean frequency (95% CI): Rad^sink^, 147.1 (145.3 – 148.8) Hz; LM^sink^, 125.4 (124.0 – 126.9) Hz; p < 10^-5^; paired bootstrap test] and lower ripple power (Figure S2H,I). These two profiles consistently emerged regardless of how the LFP and the CSD signals were referenced to ripple events (Figure S3A-E).

Next, we evaluated whether the LFP signal recorded in the pyramidal cell layer during each ripple could be predicted by the underlying CSD signals. While the CSD from any CA1 layer predicted the *stratum pyramidale* LFP waveform of Rad^sink^ and LM^sink^ ripples, the *lacunosum-moleculare* CSD remained the strongest predictor (Figures 1F and S3F). A linear discriminant classifier trained on *stratum pyramidale* LFP waveforms distinguished individual Rad^sink^ and LM^sink^ events [Figures 1G-I and S3G-K; mean accuracy gain over shuffle controls (95% CI): 55 (38.5 – 66.2) %; p = 0.0014, 1-tailed t-test; model cross-validated over 5 mice]. The pyramidal LFP waveform thus contained significant information about single-ripple CSD profiles (Figure 1I). These results demonstrate that hippocampal ripples vary in their laminar currents, with event-by-event analysis revealing a continuum that includes ripples with prominent *lacunosum-moleculare* sinks, which are not evident in the grand average profile. Moreover, the pyramidal layer LFP waveform allows distinguishing Rad^sink^ versus LM^sink^ ripples.

### Rad^sink^ versus LM^sink^ ripples exhibit distinct population-level activity patterns

We next tested whether Rad^sink^ and LM^sink^ ripples differentially engaged neuronal spiking, using multichannel tetrode recordings simultaneously from CA1 and CA3 pyramidal layers [Figure 2A; mean principal cells per recording day (IQR): 42.9 (27.0 – 62.8); 3,521 total principal cells; 280 total hours of sleep/rest from 13 mice; 244 sleep/rest sessions; mean duration (IQR): 68.8 (48.0 – 90.2) minutes per sleep/rest session]. CA3 cells are the main upstream trigger for CA1 SWRs ^4^. Using the CSD-validated model from our laminar recordings (Figure 1G-I), we identified tetrode-recorded CA1 ripples as Rad^sink^ or LM^sink^ events [Figure 2A; mean number of classified ripples per sleep/rest session IQR): Rad^sink^, 1,062 (677.8 – 1,417.0); LM^sink^, 866.8 (490.8 – 1,146.3)]. Both ripple frequency and LFP waveform of tetrode-recorded Rad^sink^ and LM^sink^ ripples (Figure S4A,B) matched those recorded with silicon probes (Figure 1D,E).

**Figure 2:**
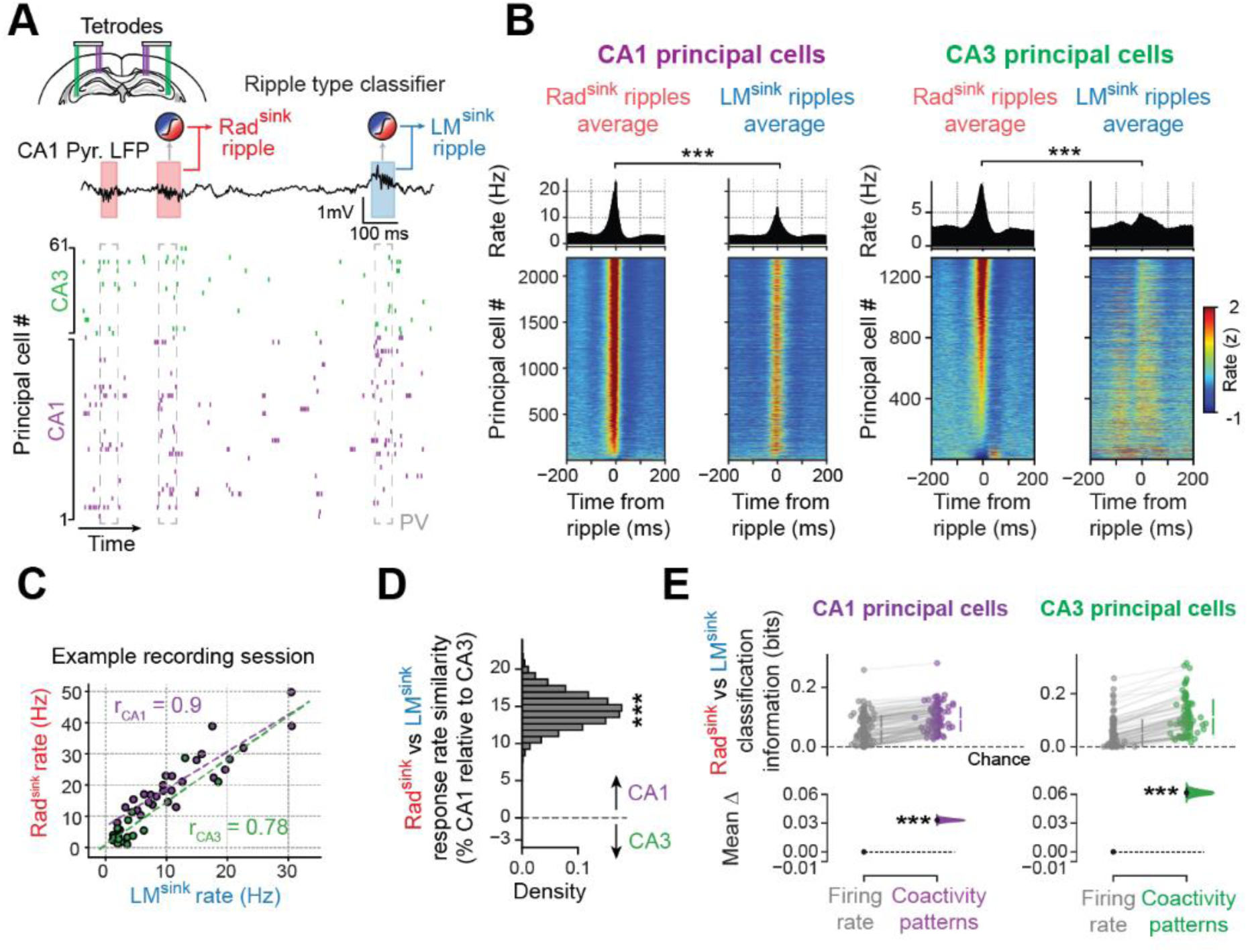
Principal cell firing response to Rad^sink^ versus LM^sink^ ripples. **(A)** Example dual-site 14-tetrode ensemble recording of CA1 and CA3 principal cells. *Top trace*, raw LFP signal from CA1 pyramidal layer. *Bottom raster plot*, (colour-coded) CA1 and CA3 principal cell spike trains (one cell per row; a 1-s sample recording for clarity). Population activity vector (PV) of individual ripple extracted using a 50-ms window centred at the ripple envelope peak. Tetrode-recorded Rad^sink^ and LM^sink^ ripples distinguished using the silicon probe-validated LDA (Figure 1G-I). **(B)** Triggered average firing response of CA1 and CA3 principal cells to Rad^sink^ and LM^sink^ ripples. *Top panels*, overall population average responses. *Bottom panels*, firing response (z-scored) of individual cells, sorted by their firing rates in Rad^sink^ ripples. **(C)** Firing response similarity of individual principal cells to Rad^sink^ versus LM^sink^ ripples from a single sleep session. Each point represents a CA1 (purple) or CA3 (green) cell. The Pearson correlation coefficient (r) measures the “response rate similarity” between Rad^sink^ and LM^sink^ ripples. Dashed coloured lines depict the corresponding best-fit linear relationships. **(D)** Mean difference in response rate similarity between CA1 and CA3 principal cells (percentage relative to rate similarity of CA3 principal cells; p < 10^-5^, bootstrap test). **(E)** Estimation plots showing the effect size for the difference in classifier accuracy (measured as mutual information) in discriminating Rad^sink^ and LM^sink^ ripples using ripple-nested PVs from CA1 (left) or CA3 (right). Each distribution is compared to chance level (dashed lines) and a surrogate distribution that preserved individual cell firing rates and overall population statistics while shuffling coactivity patterns (gray distributions). Each dot represents a single sleep session. For both CA1 and CA3, preserving population coactivity improved accuracy over both controls (CA1, p < 10^-5^; CA3, p < 10^-5^; paired bootstrap tests). *p < 0.05, **p < 0.01, ***p<0.001.

CA1 principal cell activity transiently increased in both Rad^sink^ and LM^sink^ ripples (Figure 2B), with the average CA1 firing response higher in Rad^sink^ ripples [Figure S4C; mean peak rate (95% CI): Rad^sink^, 21.88 (21.27 – 22.50) Hz; LM^sink^, 13.12 (12.70 – 13.55) Hz; p < 10^-5^, paired bootstrap test; n = 2,196 CA1 principal cells]. CA1 principal cells fired at an earlier phase in LM^sink^ ripples yet maintaining similar coupling strength (Figure S4D,E). In CA3, principal cells had higher firing rates in Rad^sink^ ripples [Figure S4F; mean peak rate (95% CI): Rad^sink^, 8.82 (8.14 – 9.54) Hz; LM^sink^, 5.18 (4.80 – 5.61) Hz; p < 10^-5^, paired bootstrap test; n = 1,325 CA3 principal cells]. CA3 principal cell firing further exhibited a distinct temporal pattern in LM^sink^ ripples, marked by a transient firing increase ∼100 milliseconds before their power peak (Figure 2B), coinciding with a radiatum sink (Figures 1B,C and S1A-C, 3A-E). Additionally, CA3 interneurons showed different Rad^sink^ versus LM^sink^ temporal firing responses (Figure S4G). In CA3, but not CA1, the ratio of interneuron over principal cell firing was higher in LM^sink^ ripples (Figure S4H). Importantly, CA1 and CA3 principal cells showed strong positive correlations in both their individual firing rates and spiking probabilities between Rad^sink^ and LM^sink^ ripples [mean Pearson correlation (95% CI): CA1, 0.86 (0.85 – 0.88); CA3, 0.75 (0.72 – 0.78)]. This high firing response similarity indicated that principal cells with strong LM^sink^ response showed strong Rad^sink^ response; cells with weak LM^sink^ response exhibited weak Rad^sink^ response (Figures 2C and S4I,J). This cross-ripple response similarity of individual principal cells was higher in CA1 than CA3 (Figures 2D and S4K).

We then tested whether Rad^sink^ and LM^sink^ ripples differed in their population-level patterns. For this, we first trained classifiers to discriminate ripple types using their instantaneous population vectors of spiking activity of either CA1 or CA3 principal cells. As controls, we used surrogate population vectors that preserved both individual neurons firing rates (i.e., number of spikes each cell discharged across ripples) and population firing rates (i.e., number of spikes the population discharged in each ripple) while shuffling the combination of neurons transiently coactive in each ripple (i.e., instantaneous population coactivity patterns). Control models outperformed chance, indicating that some information about ripple types derived from firing rates, consistent with the cross-ripple firing difference (Figure 2B and S4C,F). CA1 and CA3 population coactivity patterns distinguished Rad^sink^ and LM^sink^ ripples, surpassing control models (Figure 2E), suggesting that these two ripple types express distinct, non-redundant coactivity motifs. Ripple classification improved when the entire spike train temporal structure around the ripple event was considered, particularly for CA3 neurons, compared to the classification based solely on firing activity at ripple power peak (Figure S4L-N and Table S2). Thus, individual neurons displayed a high cross-ripple firing response similarity, but the population and temporal patterns of transiently coactive neurons differed between Rad^sink^ and LM^sink^ ripples.

### Distinct coactivity level and structure feature LM^sink^ versus Rad^sink^ population patterns

We characterised the coactivity motifs nested in Rad^sink^ versus LM^sink^ ripples. For each ripple type, we measured the coactivity of each cell pair (*i*, *j*) using the regression coefficient from the prediction of spike discharge in neuron *i* by that in neuron *j*, controlling for the remaining population (Figure 3A). Coactivity among CA1 neurons was higher in LM^sink^ ripples (Figures 3B and S4O; p < 10^-5^, paired bootstrap test), whereas CA3 neurons showed higher coactivity in Rad^sink^ ripples (Figure S4O,P).

**Figure 3:**
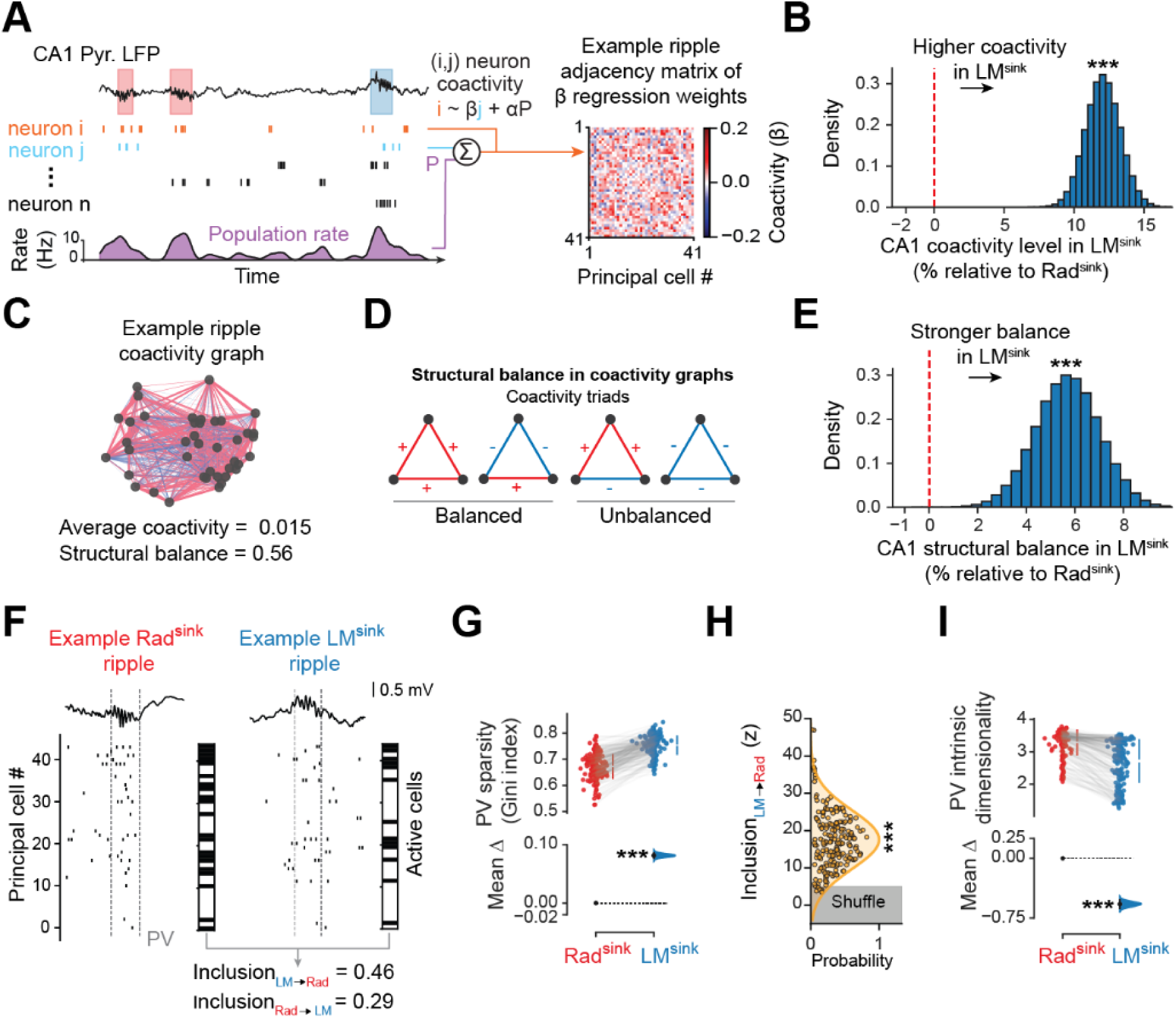
CA1 population coactivity level and structural organisation differ in Rad^sink^ versus LM^sink^ ripples. **(A)** Schematic outlining the measure of ripple-nested principal cell coactivity. *Left*, an example LFP trace from the CA1 pyramidal layer accompanies a raster plot of a subset of CA1 principal cells. Neuron *i* spiking is predicted by neuron *j* spiking while accounting for the population rate (*P*) using a linear regression model. *Right*, corresponding ripple adjacency matrix of beta coefficients. **(B)** Mean paired difference reveals that CA1 principal cells exhibit higher coactivity in LM^sink^ compared to Rad^sink^ ripples. Difference expressed as a percentage relative to the mean coactivity in Rad^sink^ ripples. **(C)** Example neuronal graph representing the ripple adjacency matrix shown in (A), including its average coactivity and structural balance. **(D)** Schematics of balanced and unbalanced triads of coactivity used to assess structural balance in ripple-nested neuronal graphs. **(E)** Mean paired difference (as in B) shows that LM^sink^ ripple coactivity graphs have higher structural balance than Rad^sink^ ripple coactivity graphs. **(F)** Raw example LFP trace and raster plot of CA1 principal cells for one Rad^sink^ ripple and one LM^sink^ ripple, with corresponding binarized population vectors (PVs) representing their cell recruitment. The overlap between these two binarized PVs defines their cross-ripple inclusion measure. **(G)** Estimation plot showing the effect size for the difference in the sparsity (Gini index) of the population vectors nested in Rad^sink^ versus LM^sink^ ripples. Each point represents the mean sparsity of a single sleep session. **(H)** Distribution of the average inclusion values of motifs of active principal cells in *LM* → *Rad* ripples. Each dot represents a sleep session. Inclusion values z-scored relative to surrogate distributions generated by shuffling cell coactivity. Gray shaded region shows ± 5 standard deviations. **(I)** Estimation plots showing the effect size for the difference in intrinsic dimensionality of population vectors in Rad^sink^ versus LM^sink^ ripples. Each point represents a single sleep session. *p < 0.05, **p < 0.01, ***p<0.001.

The combination of positively and negatively correlated spike trains shapes the stability of the underlying network of firing relationships ^32^. We thus determined the stability of the population patterns in LM^sink^ and Rad^sink^ ripples by computing the structural balance of the neuronal graphs embedding the measured coactivity relationships (Figure 3A). In these graphs, each node represents one cell, and each edge represents the coactivity of one cell pair (Figure 3C). We then detected balanced and unbalanced motifs of (triadic) relationships (Figure 3D). LM^sink^ ripples contained a stronger proportion of balanced coactivity motifs in the CA1 population (Figure 3E; p = 6×10^-5^, paired bootstrap test). Structural stability in the CA3 population activity was similar across ripple types (Figure S4Q). These results indicated that LM^sink^ ripples contain more stable motifs of CA1 coactivity.

We noted that CA1 population vectors appeared sparser in LM^sink^ ripples (Figure 3F,G; Gini index; p < 10^-5^, bootstrap test), recruiting fewer CA1 principal cells at a time [Figure S5A; mean proportion of active CA1 principal cells (95% CI): Rad^sink^, 41.4 (40.5 – 42.3) %; LM^sink^, 28.0 (27.2 – 28.7) %; p < 10^-5^, paired bootstrap test]. This reduced recruitment of CA1 principal cells indicated expression of smaller motifs of stronger coactivity in LM^sink^ ripples. Transiently recruited CA1 principal cells yet responded similarly across ripple types (Figures 2C,D and S4I-K).We thus explored whether such core motifs could reoccur within the larger (denser) motifs of Rad^sink^ ripples. We used an asymmetrical Jaccard coefficient to measure the tendency for sets of neurons coactive in LM^sink^ ripples to be contained within the coactive population observed in Rad^sink^ ripples. This directional procedure quantified, for each pair of ripples (*m*, *n*), the extent to which the set of active cells in ripple *m* was also active in ripple *n* (Figures 3F and S5B,C). Coactivity motifs forming LM^sink^ population vectors showed higher representation in Rad^sink^ vectors than a control distribution obtained by shuffling cells jointly active in each LM^sink^ ripple [Figures 3H; mean inclusion *LM*→*Rad* (95% CI): z-scored against controls: 19.53 (18.37 – 20.70); p = 5×10^-56^, 1-tailed t-test].

The LM^sink^ motifs of coactive neurons can therefore appear in Rad^sink^ ripples alongside additional cells. This suggests that LM^sink^ ripples host generic (lower-dimensional) population patterns with core motifs of coactivity, while Rad^sink^ ripples nest composite (higher-dimensional) patterns that incorporate those from LM^sink^. Accordingly, LM^sink^ population vectors exhibited lower intrinsic dimensionality ^33^ [Figure 3I; mean intrinsic dimensionality (95% CI): Rad^sink^, 3.29 (3.24 – 3.35); LM^sink^, 2.72 (2.63 – 2.81); p < 10^-5^, paired bootstrap test]. Thus, LM^sink^ and Rad^sink^ population patterns differ in coactivity level and structure, with the lower-dimensional neuronal motifs featured in LM^sink^ ripples re-appearing within Rad^sink^ ripple population vectors.

### Neuronal content of CA1 population patterns in Rad^sink^ and LM^sink^ ripples

Traditionally, the network operations supporting hippocampal functions have been studied under the assumption that each region (e.g., CA1) contains a homogeneous population of functionally equivalent principal cells. Recent research has now revealed significant heterogeneity within hippocampal principal cell populations, segregating CA1 principal cells into two parallel channels with distinct properties serving differently information processing ^34–38^. CA1 principal cells with somatic location in the deep pyramidal sublayer exhibit higher rate and more rigid activity; their superficial counterparts show lower firing rate and more plastic recruitment ^32,39–50^. We thus assessed whether LM^sink^ versus Rad^sink^ ripples engaged differently deep and superficial cells recorded along the radial axis of the CA1 pyramidal layer (Figure 4A) ^51^. For both populations of principal cells, we calculated the change in their firing rates from pre-ripple baseline to each ripple type, accounting for differences in individual average rates. During Rad^sink^ ripples, superficial and deep cells increased their firing similarly (Figure 4B,C); whereas during LM^sink^ ripples, deep cells showed higher increase in firing rates than superficial cells, which indicated stronger recruitment of deep cells in LM^sink^ ripples. Additional analysis using conditional firing probabilities confirmed preferential recruitment of superficial cells in Rad^sink^ but not LM^sink^ ripples (Figure 4D).

**Figure 4:**
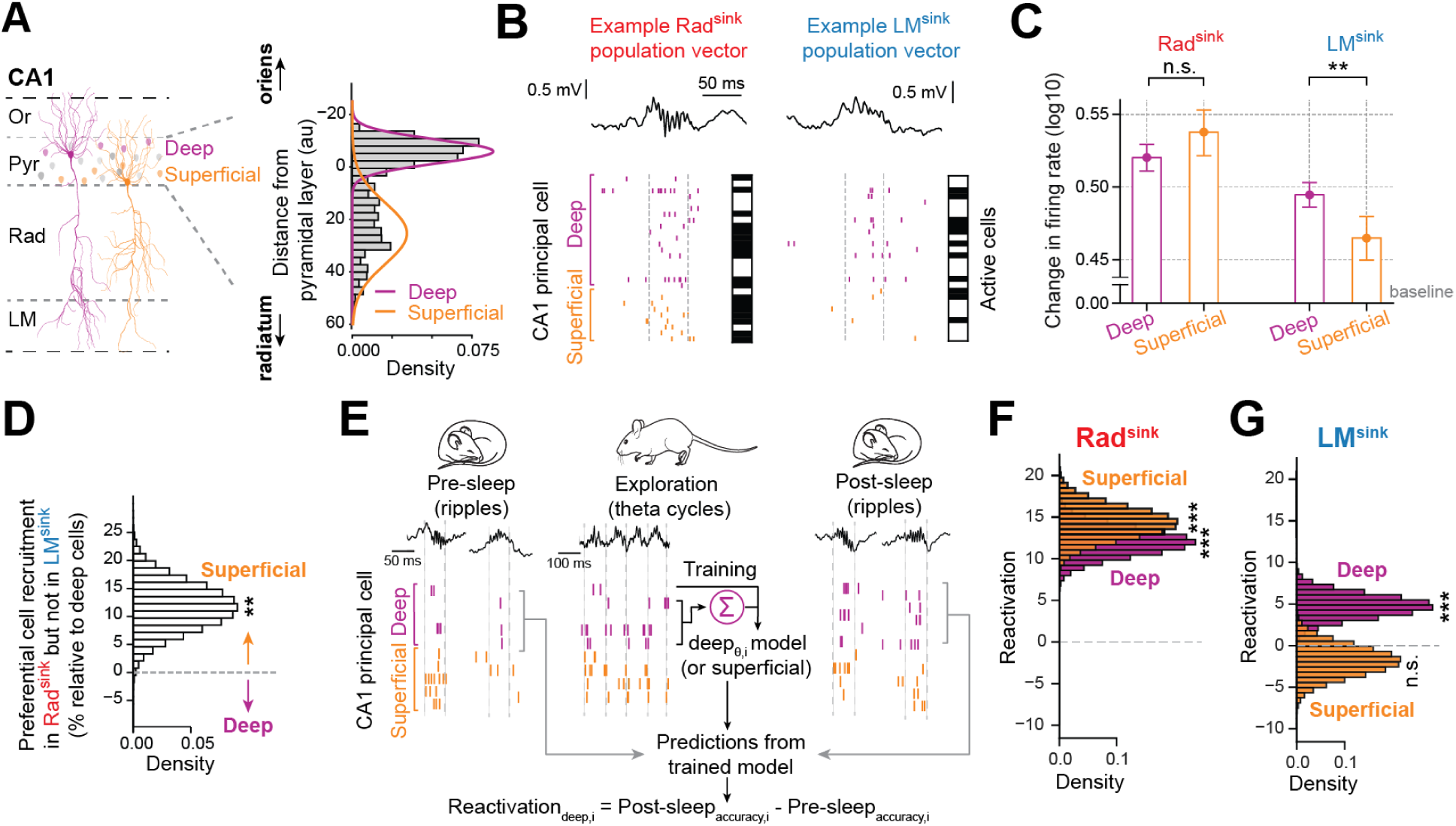
Firing response to Rad^sink^ and LM^sink^ ripples of principal cells in deep versus superficial CA1 pyramidal sublayers. **(A)** Distribution of CA1 principal cells recorded in the deep (purple) versus superficial (orange) pyramidal sublayers (n = total 1,353 deep and 843 superficial cells). **(B)** Example raw LFP traces and raster plots showing (colour-coded) spiking activity of deep and superficial cells in Rad^sink^ and LM^sink^ ripples, with their corresponding binary population vectors depicting recruited cells. **(C)** Change in firing response of deep and superficial cells. In Rad^sink^ ripples, deep and superficial principal cells increase their firing rates similarly (relative to their pre-ripple baseline), whereas in LM^sink^ ripples, deep cells show a significantly greater firing increase compared to superficial cell (mean ± 95% CI; Rad^sink^, p = 0.10; LM^sink^, p = 3.2×10^-3^; bootstrap tests). **(D)** Preferential cell recruitment in Rad^sink^ ripples. We computed the conditional probability of cells to be selectively active in Rad^sink^ ripples (given their inactivity in LM^sink^ ripples), normalised relative to chance, and shown as a percentage relative to the mean of deep cells. Superficial cells were more likely than deep cells to be active in Rad^sink^ and not in LM^sink^ ripples (p = 2.7×10^-3^; bootstrap test). *Dashed line*, chance level **(E-G)** Rad^sink^ ripples reactivate waking theta coactivity of both deep and superficial cells but LM^sink^ ripples selectively reactivate deep cells. Shown in (E) is a schematic of the method for computing offline reactivation. Using theta cycles in active exploration, a generalized linear model (GLM) was trained to predict the activity of each principal cell from the waking theta spiking activity of the other cells in the same sublayer. Each theta model was then applied to predict the response of its target cell during Rad^sink^ and LM^sink^ ripples of pre-exploration sleep versus post-exploration sleep. Reactivation was measured as the difference in model accuracy in post-exploration sleep relative to pre-exploration sleep. Shown in (F) is the mean difference in reactivation during Rad^sink^ ripples for deep and superficial cells. Both sublayers exhibit significant Rad^sink^ reactivation [all Ps < 10^-5^; 1-tailed bootstrap tests]. (G) as (F), but for LM^sink^ ripples. Note that superficial cells do not reactivate significantly during post-sleep LM^sink^ ripples [deep cells: p = 2.7×10^-4^; superficial cells: p = 0.84; 1-tailed bootstrap tests]. *p < 0.05, **p < 0.01, ***p < 0.001.

Hippocampal ripples reactivate offline the neural patterns expressed in previous wakefulness^4^. To examine the contribution of deep and superficial cells in offline reactivation during Rad^sink^ and LM^sink^ ripples, we trained generalized linear models to predict the spiking activity of each principal cell based on its sublayer peers during theta oscillations of exploratory behaviour (Figure 4E; a leave-one-out approach where deep cells predict another deep cell; superficial cells predict another superficial cell). We then measured reactivation by applying the theta-nested coactivity model of each cell to predict its ripple-nested activity in post-exploration sleep/rest, controlled for pre-exploration sleep/rest. We found that Rad^sink^ ripples reactivated the waking theta activity of both deep and superficial cells, whereas LM^sink^ ripples selectively reactivated deep cells (Figures 4F,G and S6A; all Ps < 10^-5^, except for superficial cells in LM^sink^ ripples: p = 0.84; 1-tailed bootstrap tests). CA3 principal cells mirrored CA1 superficial cells, showing significant reactivation in Rad^sink^ but not in LM^sink^ ripples (Figure S6B).

### Temporal stability of Rad^sink^ recent patterns and gradual drift of LM^sink^ prior patterns

These findings show that the population response to Rad^sink^ and LM^sink^ ripples exhibit important differences with respect to activity level, structure, and neuronal content. What could be the significance of such differences? Rad^sink^ ripples contain higher-dimensional population patterns reactivating superficial cells combined with deep cells into dense coactivity motifs. LM^sink^ ripples nest lower-dimensional population patterns that primarily engage deep cells in core motifs of coactivity. We thus explored how these differences in the population response to Rad^sink^ versus LM^sink^ ripples reflect the population spiking patterns seen during wakefulness.

We decomposed neuronal coactivity motifs into ‘recent’ motifs (i.e., neurons selectively coactive during arena exploration but not during the preceding sleep/rest) and ‘prior’ motifs (i.e., neurons already coactive in pre-exploration sleep/rest). We then measured the reactivation strength of these “recent” versus “prior” motifs during post-exploration sleep/rest (Figures 5A and S6C-F). Strikingly, Rad^sink^ ripples reactivated recent coactivity motifs while LM^sink^ ripples continued to express prior motifs (Figure 5B; prior motifs: p = 6×10^-4^; recent motifs: p < 10^-5^; bootstrap tests). LM^sink^ coactivity motifs indeed showed strong cross-sleep similarity (Figure S6G), consistent with stronger structural balance (Figure 3E) and lower dimensionality of LM^sink^ patterns (Figure 3I). Deep cells predominantly contributed to prior coactivity motifs (Figure 5C; p = 2×10^-5^; bootstrap test), while superficial cells contributed more significantly to the reactivation of recent coactivity motifs (Figure 5D; p = 1.2×10^-4^, bootstrap test). LM^sink^ ripples thus host core motifs of prior neuronal coactivity that engage deep cells and are stable across sleep/rest epochs. Rad^sink^ ripples reactivate recent motifs of waking coactivity that recruit superficial cells, combining them with deep cells into high-dimensional population patterns.

**Figure 5:**
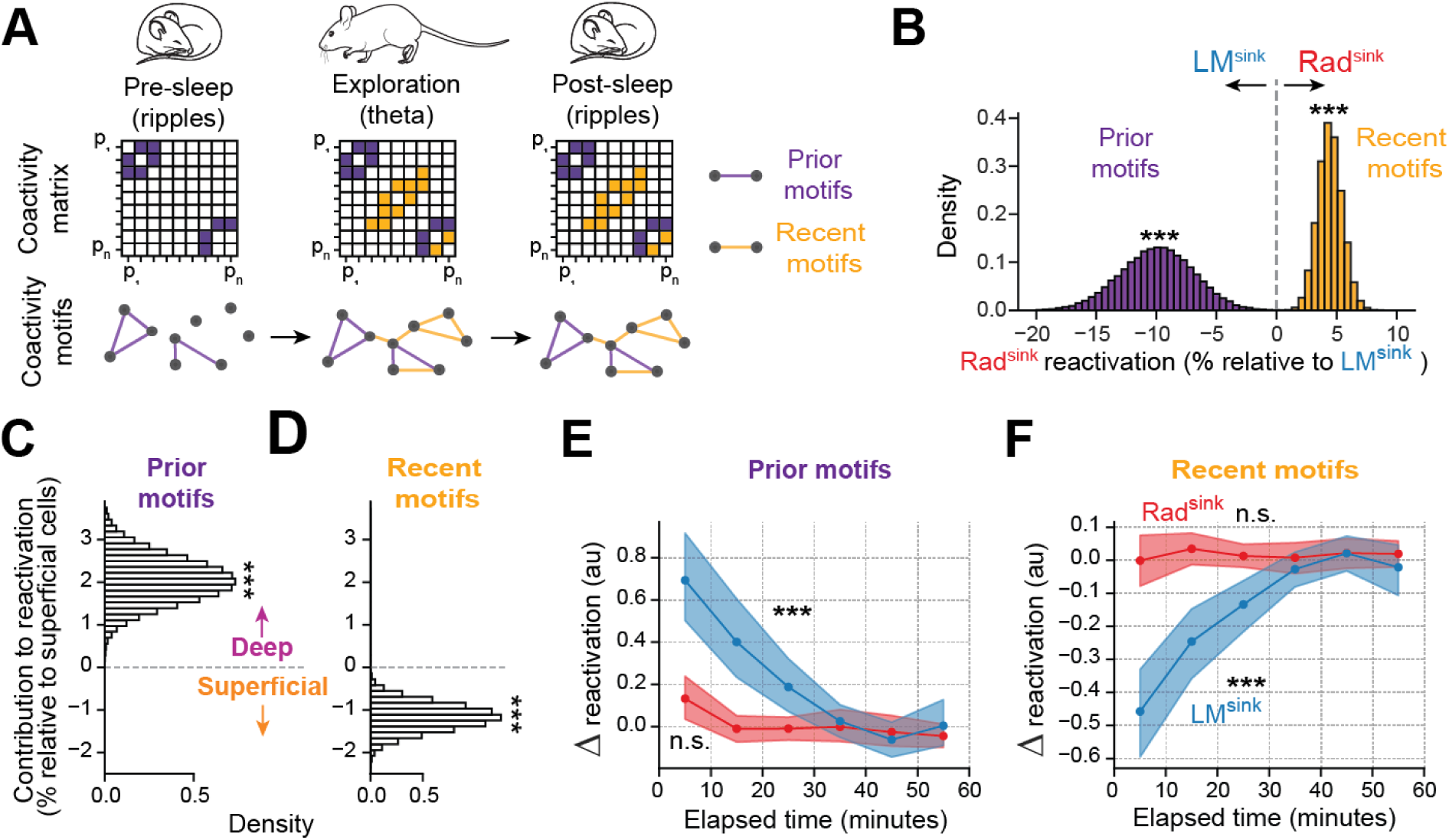
Rad^sink^ and LM^sink^ reactivation dynamics of prior versus recent coactivity motifs. **(A)** Schematic of the method identifying prior and recent motifs of neuronal coactivity (see also Figure S6C-F). Coactivity matrices (top) and corresponding neuronal graph motifs (bottom) across pre-sleep (ripples), exploration (theta cycles), and post-sleep (ripples). Purple squares denote cell pairs constituting prior coactivity motifs (i.e., already present in pre-sleep), while orange squares show coactivity relationships that selectively emerged during exploration (i.e., absent in pre-sleep). **(B)** Mean difference in reactivation during Rad^sink^ post-sleep ripples (relative to LM^sink^ ripples) for prior and recent motifs. Rad^sink^ ripples selectively reactivated recently expressed theta coactivity motifs, whereas LM^sink^ ripples preferentially reactivated prior motifs **(C,D)** Mean difference in the contribution of deep and superficial cells to the reactivation of prior (C) and recent (D) motifs (relative to the mean of superficial cells). Deep cells primarily contributed to prior motifs, whereas superficial cells predominantly contributed to recent motifs **(E,F)** Time course for the mean change in reactivation of prior (E) versus recent (F) motifs in Rad^sink^ (red) and LM^sink^ (blue) ripples over the hour-long sleep/rest (normalized by subtracting the overall mean reactivation of that session). LM^sink^ reactivation of prior motifs showed an exponential decay while that of recent motifs showed an exponential increase. Rad^sink^ ripples did not exhibit such temporal changes, allowing temporally stable reactivation throughout hour-long sleep/rest. Significance levels determined by the fitness score of the exponential function. *Shaded areas*, 95% CI. *p < 0.05, **p < 0.01, ***p < 0.001.

LM^sink^ ripples nest strong coactivity among deep cells, but can plastic changes nevertheless occur in such core motifs over longer timescales? We analysed temporal dynamics of CA1 population patterns during extended periods of sleep [mean duration (IQR): 86.25 (76.33 – 91.62) minutes per sleep]. Strikingly, in LM^sink^ ripples, the strength of prior motifs (i.e., those already present in pre-exploration sleep) decreased exponentially during the first half hour of post-exploration sleep/rest [Figure 5E; time constant *τ* (95% CI), 13.12 (8.34 – 21.80) minutes; p < 10^-4^; 1-tailed bootstrap test] while the strength of recent motifs increased exponentially over the same period [Figure 5F; time constant *τ* (95% CI): 13.77 (8.50 – 27.70) minutes; p < 10^-4^; 1-tailed bootstrap test]. Rad^sink^ ripples showed no such changes in their reactivation content throughout the hour-long sleep/rest [Figure 5E,F; prior motifs, p = 0.32; recent motifs, p = 0.23; bootstrap tests]. Changes in LM^sink^ reactivated content, from prior to recent motifs, over the hour-long post-exploration sleep/rest, were explained by ripple occurrence time and not by ripple occurrence frequency nor ripple population sparsity (Figure S6H-J). Following wakefulness, the prior coactivity motifs expressed in LM^sink^ ripples thus gradually drifted over time towards the recent waking activity subspace while maintaining a consistent proportion of active cells.

## Discussion

In this study, we identified two ripple profiles based on their distinct laminar currents across the CA1 somato-dendritic axis, that differentially recruited hippocampal neurons during offline periods of sleep/rest. The first profile, with stronger radiatum currents, integrates superficial cells with deep CA1 principal cells into higher-dimensional composite patterns that undergo hour-long stable reactivation of recently expressed waking motifs of neuronal coactivity. The second ripple profile, with stronger lacunosum-moleculare currents, contains core motifs of deep cells forming lower-dimensional patterns that undergo time-varying reactivation, gradually drifting over sleep from prior to recent coactivity spaces. Collectively, these findings reveal a diversity in hippocampal ripple profiles linked to the offline organisation of neuronal population reactivation. By tuning differences in the activity level, structural organisation, and neuronal content of population activity patterns (Figure 6), this ripple-by-ripple diversity could instantiate parallel reactivation channels for stable integration of recent wakefulness and flexible updating of prior internal representations.

**Figure 6:**
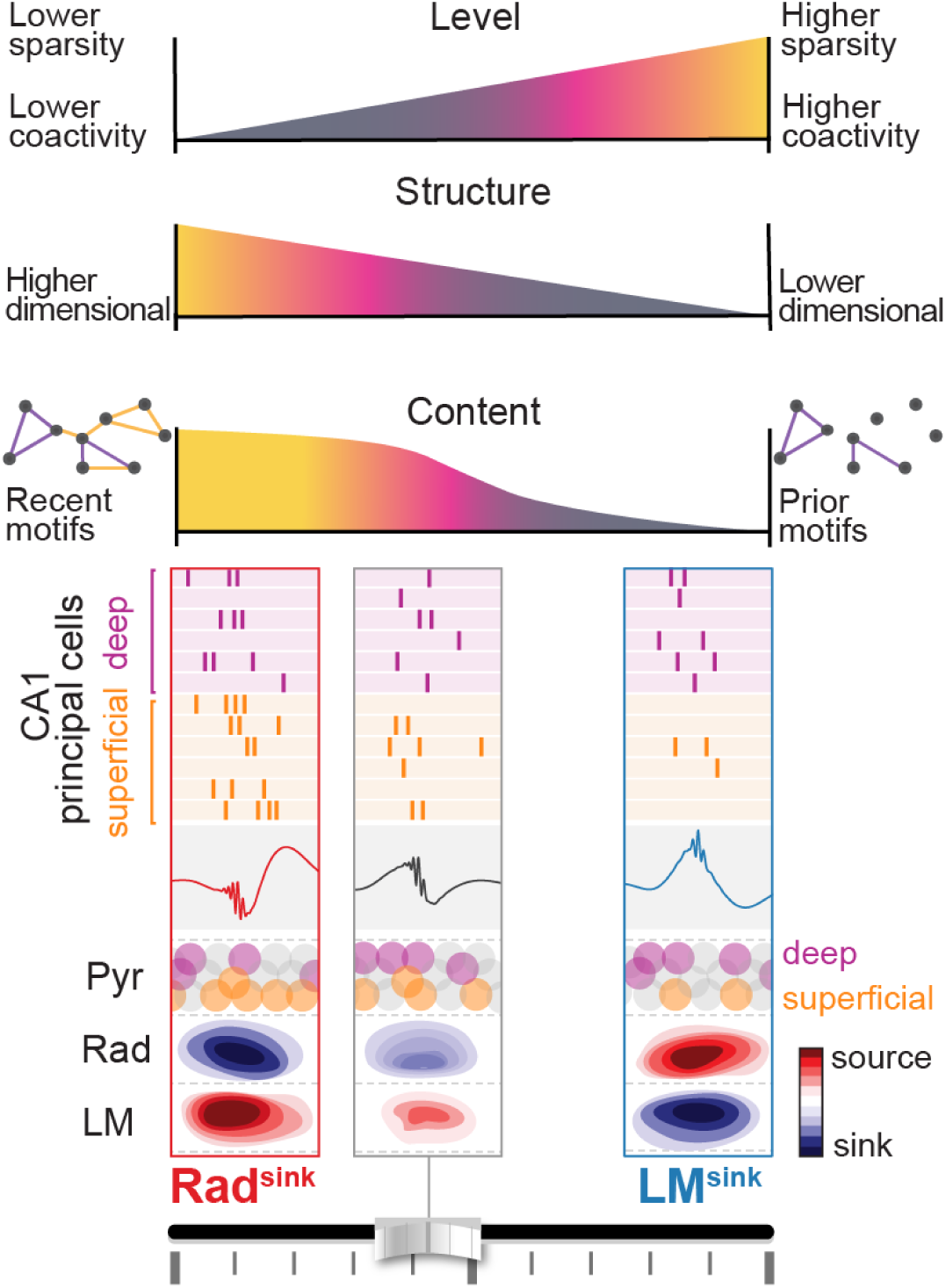
Summary schematic. Two ripple profiles, identified by their radially organised currents, exhibit distinct CA1 population activity level, structure, and content. Rad^sink^ ripples, with stronger *radiatum* current sinks, integrate recent motifs of waking coactivity, combining superficial with deep principal cells into higher-dimensional population patterns that undergo stable reactivation over hour-long sleep/rest. In contrast, LM^sink^ ripples, with stronger *lacunosum-moleculare* current sinks, contain core motifs of pre-structured coactivity, engaging deep cells into lower-dimensional patterns that undergo a reactivation drift, gradually updating prior for recent activity content.

We began our investigation by analysing the variability in CSD profiles of individual ripple events. These profiles form a continuum rather than discrete clusters, aligning with reports suggesting that ripple waveforms and features occupy a continuum of parameter-space ^28^. This might enable the dynamical modulation of the hippocampal population activity on a ripple-by-ripple basis using transient changes of currents in specific neural layers. The dominant contributors to this continuum are ripples associated with a stronger current sink in *stratum radiatum* (Rad^sink^ ripples), reflecting the canonical CSD profile of the grand average ripple. Interposed with these are ripples that instead show a stronger current sink in *stratum lacunosum-moleculare* (LM^sink^ ripples), thus far hidden in the grand average. CA3 projections to CA1 *stratum radiatum* play a key role in sharp-wave/ripple generation ^4,52,53^, consistent with the observed large average input current in this layer (Figure 1A). Yet, both this study and previous work ^28^ reveal significant variability in current profiles associated with individual ripples. Using an unsupervised approach, we identified a subset of ripples with a stronger current sink in *stratum lacunosum-moleculare* and slower ripple frequency. This LM^sink^ ripple profile likely implicates projections from the medial entorhinal cortex layer III (EC3) to CA1, suggesting a complex interplay of CA3 and EC3 inputs in CA1 ripple generation. In line with this, removing CA3 inputs does not entirely prevent ripple generation; instead, slower-frequency ripples persist in the absence of CA3 ^53^. Furthermore, recent findings indicate that EC inputs to CA1 can shape ripples characteristics ^49,54,55^. This transient tuning on an moment-by-moment basis across individual ripples is reminiscent of that previously observed across individual cycles of hippocampal theta oscillations during awake exploration ^46,56^. Similar to waking theta cycles, our findings show that the hippocampus dynamically tunes offline population activity on a ripple-by-ripple basis, in line with recent work suggesting that a temporal microstructure of sleep embeds the reactivation of co-existing hippocampal patterns ^57^.

We found that principal cells and interneurons in both CA1 and CA3 were recruited at varying levels across the two ripple profiles. Rad^sink^ ripples have higher firing rates and denser recruitment of neurons compared to LM^sink^ripples. Furthermore, in CA3, beyond these quantitative differences in neuronal recruitment, we observed qualitatively distinct temporal firing responses in Rad^sink^and LM^sink^ ripples. These findings further support the idea that these two ripple-associated CSD profiles arise in the hippocampal circuit differently. The high recurrency in CA3 could enable strong excitatory currents converging to *stratum radiatum* during Rad^sink^ ripples, activating a larger number of neurons at higher firing rates and producing higher-frequency ripples. In contrast, during LM^sink^ ripples, CA3→CA1 inputs would be significantly weaker, either intrinsically or attenuated locally by interneurons within CA3 or CA1, promoting stronger differential current in *stratum lacunosum-moleculare*. These distal currents still recruit a notable population of neurons but at lower firing rates than in Rad^sink^ ripples. This reduced recruitment may stem from weaker input currents from CA3 during LM^sink^ ripples or from the additional attenuation caused by the longer distance these currents must travel to generate somatic spikes in CA1, compared to the shorter path from *stratum radiatum*. Regardless, our findings establish that within the continuum of CSD profiles, CA1 and CA3 principal cells are modulated to varying degrees, reinforcing the dynamic nature of ripple-associated recruitment in the hippocampal circuitry.

The differences between Rad^sink^ and LM^sink^ ripples extend beyond modulation of activity levels to include distinct structural properties of the population patterns formed by principal cell coactivation in these events. Specifically, we show that different coactivity motifs are recruited across the two ripple profiles. CA1 principal cells contributing to the sparser LM^sink^ ripples formed core motifs of high and robust coactivity, which can also reappear in Rad^sink^ ripples but alongside additional cells. In this manner, whilst LM^sink^ ripples nest lower-dimensional patterns, Rad^sink^ ripples nest composite, higher-dimensional population patterns. These findings suggest that, despite potentially originating from parallel hippocampal circuits, CA1 principal cell population responses to Rad^sink^ and LM^sink^ ripples can influence each other.

Recent studies have documented significant diversity among CA1 pyramidal cells, emphasizing that their somatic location within the deep versus superficial sublayers of the *stratum pyramidale* predicts distinct contributions to hippocampal network dynamics ^32,40,58–60^. Overall, during ripples, superficial CA1 principal cells exhibit stronger changes in firing activity compared to deep cells ^32,40,51,58^. Here we show that while Rad^sink^ ripples reflect the canonical pattern of superficial cells being more active than deep cells, LM^sink^ ripples exhibit the opposite trend, with deep cells being predominantly more active than their superficial counterparts. Furthermore, we found that the additional CA1 principal cells recruited during Rad^sink^ ripples – but not during LM^sink^ ripples – are primarily superficial cells. Importantly, while previous studies showed that the offline reactivation of waking patterns in CA1 is mainly driven by superficial cells ^58^, we observed that this is not the case during LM^sink^ ripples. CA1 superficial cells and CA3 principal cells were not significantly reactivated during these events. These findings align with earlier reports showing that CA1 deep cells receive stronger inputs from EC3, which projects to *stratum lacunosum-moleculare*; while CA1 superficial cells receive stronger inputs from CA3, which projects to *stratum radiatum* ^37,40,45,61^. Accordingly, we suggest that during Rad^sink^ ripples, offline reactivation of waking patterns primarily engages CA3 ensembles, which in turn bias CA1 to reactivate the associated higher-dimensional response integrating superficial CA1 cells.

Deep and superficial CA1 principal cells have also been shown to respond differently to behavioural experiences. Notably, in theta cycles marking wakeful exploration, deep cells exhibit higher firing rates and more stable activity patterns, whereas superficial cells show lower firing rates and adapt flexibly to new experiences ^32,51,58^. This difference could be partly attributed to stronger excitatory inputs that superficial cells receive from CA3, enabling them to respond to contextual changes ^37,40,48^. Consistent with these findings, we show that superficial cells drive in Rad^sink^ ripples of post-exploration sleep the expression of coactivity motifs expressed during the preceding exploratory behaviour. In contrast, the more rigid deep cells instantiate pre-existing motifs of coactivity in LM^sink^ ripples. Previous studies reported that CA1 superficial, but not deep, principal cells undergo postsynaptic potentiation in response to CA3 inputs after novel experience ^48^. These results suggest that CA3 enables CA1 superficial cells to rapidly and flexibly reorganise coactivity patterns in Rad^sink^ ripples following new waking experience. Conversely, the reduced activity of CA3 during LM^sink^ ripples could favour the expression of deep cell motifs.

Why does the hippocampus co-process offline rapid integration and gradual stabilisation of recent and prior population responses, respectively? While the prompt integration of new information is essential, maintaining a stable underlying population activity structure could yet provide a “backbone” upon which finer-grain information processing can be built ^62^. Recent theoretical work suggests that the hippocampus integrates new sensory information using a pre-structured internal scaffold provided by EC, enabling scalable, flexible, and efficient memory storage ^63^. In line with this, the core motifs of deep cells associated here with LM^sink^ ripples would define such a pre-structured scaffold. Recent waking experience would then flexibly recruit coactivity motifs of CA1 superficial cells within the network. Through CA3-driven activity, the network is transiently pushed out of the prior neural backbone to append new information. Combining deep and superficial cells would allow enhancing the specificity and distinctiveness of newly-acquired representations ^62^. The resulting composite population patterns would be later reactivated offline during Rad^sink^ ripples. Meanwhile, LM^sink^ ripples would continue expressing core activity motifs, which gradually drift to update the population backbone. Deep and superficial cells would therefore work synergistically towards the concomitant rapid integration and gradual refinement of hippocampal population representations.

The prior coactivity motifs nested in LM^sink^ ripples are more stable and yet not stationary over the hour-long sleep following recent wakefulness. Population patterns in LM^sink^ ripples, but not those in Rad^sink^ ripples, gradually drift toward the coactivity space marking the recent waking activity. It remains for future studies to reveal the mechanistic implementation of this drift, occurring while the most recently activated coactivity motifs are quickly integrated into high-dimensional patterns of deep and superficial cells in Rad^sink^ ripples. An interesting avenue concerns the input-output relationships established by superficial and deep CA1 principal cells, and the possibility of bidirectional influences. Indeed, CA1 superficial cells predominantly project to EC ^36,37,45,58,61^; CA1 deep cells preferentially receive inputs from EC. This could actuate a feedback loop to CA1 deep cells. CA1 superficial cells could also modulate deep cells by recruiting local inhibitory circuits (e.g., parvalbumin-expressing basket cells ^40,64^). By these means, CA1 superficial cells would influence both the hippocampal-entorhinal and the intra-hippocampal circuitries, shaping the core motifs of deep CA1 cell activity embedded in LM^sink^ ripples. That is, during post-exploration sleep, motifs of superficial cells reflecting recent waking experience reactivate in Rad^sink^ ripples. This could in turn bias EC activity, which projects back to CA1 deep cells, gradually shifting the CA1 deep motifs toward the most recent activity state space. This interactive process could help internal linking of the recent experience with previous ones in the hippocampus, and allow network updating with the most recently encountered information ^65,66^. Moreover, CA1 deep cells establish important projections to neocortical areas (prefrontal cortex) ^58^. The gradual drift of the hippocampal backbone, evolving from prior to recent coactivity spaces over hour-long post-exploration sleep/rest, could exert important influence on systems consolidation of neuronal ensembles in downstream neocortical circuits (e.g., prefrontal cortex ^67^). In parallel, this gradual drift would also update the local neuronal priors within the hippocampus, which would then serve as the foundation for the next low-dimensional backbone in LM^sink^ ripples, while Rad^sink^ ripples continue to push the network out of this state through superficial cells, with high-dimensional patterns reporting recent wakefulness. Such a functional loop could support a network trade-off between stability and flexibility.

These findings underscore the contribution of the diversity in the expression profiles of a given network pattern (e.g., CA1 ripples), which along the heterogeneity within a given neuronal population (e.g., CA1 principal cells), can instantiate parallel processing channels in the hippocampus. Here, Rad^sink^ ripples nest hour-long, temporally stable reactivation of recent waking experience, integrating recently recruited superficial cells with core activity motifs of deep cells. In contrast, LM^sink^ ripples nest time-varying reactivation of prior population patterns, undergoing hour-long gradual drift that updates core activity motifs with recent waking experience. This hippocampal backbone could not only preserve a coherent population activity structure but also facilitates automatic integration of new waking information. Organising distinct waking experiences within a common “schema” could support flexible computations and further reduce energy demand compared to a framework imposing the *de novo* creation of entirely new high-dimensional population patterns from the outset each time ^63,68–70^. We propose that ripple-by-ripple diversity leverages differences in the activity level, structural organisation, and neuronal content of population patterns for parallel offline reactivation of prior versus recent activity in the hippocampal system.

## Acknowledgements

This work was supported by the Medical Research Council (MRC) UK (Programmes MC_UU_12024/3, MC_UU_00003/4, and MR/W004860/1 to D.D.) and the Engineering and Physical Sciences Research Council (EPSRC) UK (Awards EP/V013068/1, EP/V03474X/1, and EP/Y028872/1 to R.L.). M.C. is supported by an MRC UK studentship (MC_ST_BNDU_2019).

## Author contributions

Conceptualization, M.C., V.L.-d.-S., and D.D.; Methodology, M.C., V.L.-d.-S., G.P.G., R.L., and D.D.; Formal Analysis, M.C.; Data Collection, V.L.-d.-S.; Investigation, M.C. and V.L.-d.-S.; Resources, M.C., V.L.-d.-S., G.P.G., and D.D.; Writing – Original Draft, M.C., V.L.-d.-S., and D.D.; Writing – Reviewing & Editing, M.C., V.L.-d.-S., G.P.G., R.L., and D.D.; Visualization, M.C., V.L.-d.-S., and D.D.; Supervision, D.D; Funding Acquisition, D.D.

## Materials and Methods

### Animals

These experiments used adult (4–6 months old) C57BL/6J wild-type mice (Charles River Laboratories, UK). Animals were housed with their littermates up until the start of the experiment. All mice were held in IVCs, with wooden chew sticks and nestlets in a dedicated housing facility with a 12/12 h light/dark cycle (lights on at 07:00), 19–23°C ambient temperature and 40–70% humidity. They had free access to water and food *ad libitum* throughout the experiment. Experimental procedures were performed on mice in accordance with the Animals (Scientific Procedures) Act, 1986 (United Kingdom), with final ethical review by the Animals in Science Regulation Unit of the UK Home Office.

### Surgical procedure

All surgical procedures were performed under deep anaesthesia using isoflurane (0.5–2%) and oxygen (2 l/min), with analgesia provided before (0.1 mg/kg vetergesic) and after (5 mg/kg metacam) surgery.

For silicon probe recordings, mice were implanted with a single-shank silicon probe (Supplemental Table S1) under stereotaxic control in reference to bregma, using central coordinates −2.0 mm anteroposterior from bregma, +1.7 mm lateral from bregma, and an initial depth of 1.5 mm ventral from the brain surface to span the somato-dendritic axis of CA1 principal cells and reach the DG. Following the implantation, the exposed parts of the silicon probe were covered with Vaseline® Healing Jelly, after which its plastic drive was secured to the skull using dental cement and stainless-steel anchor screws inserted into the skull. Two of the anchor screws, both above the cerebellum, were attached to a 50 µm tungsten wire (California Fine Wire) and served as ground. For the recordings, the silicon probe was positioned along the radial axis of CA1 pyramidal cells, using the rotations applied to its holding screw.

For tetrode recordings, mice were similarly implanted with a single microdrive containing 14 independently movable tetrodes, each positioned to target the *stratum pyramidale* of either CA1 or CA3 in the dorsal hippocampus. Tetrodes were constructed by twisting together four insulated tungsten wires (12 μm diameter, California Fine Wire) which were briefly heated to bind them together into a single bundle. Each tetrode was loaded in one cannula attached to a 6 mm long M1.0 screw to enable its independent manipulation of depth. The drive was implanted under stereotaxic control in reference to bregma using the following coordinates. For CA1 pyramidal cell layer tetrodes, the span was between AP −2.0 to –2.4 mm and ML 1.6 to 2.3 mm. For CA3 pyramidal cell layer tetrodes, the span was between AP −1.8 to –2.2 mm and ML 2.0 to 2.7 mm. The initial depth of the tetrodes during the implantation surgery was 1.0 mm ventral from the brain surface. The distance between neighbouring tetrodes was 350 μm. Following the implantation, the exposed parts of the tetrodes were covered with paraffin wax, after which the drive was secured to the skull using dental cement and stainless-steel anchor screws inserted into the skull. Two of the anchor screws, both above the cerebellum, were attached to a 50 µm tungsten wire (California Fine Wire) and served as ground. For the recordings, each tetrode was lowered along the vertical axis to reach either the CA1 or CA3 pyramidale layers, using the rotations applied to its tetrode cannula-holding screw and the electrophysiological profile of the local field potentials in the hippocampal ripple frequency band, with final depth position subsequently confirmed by histology of anatomical tracks.

### Recording Procedure

Following implantation surgery, mice were allowed to recover for at least seven days. They were then familiarized with the recording procedure, being handled daily in a dedicated towel, connected to the recording cable and exposed to the sleep box for at least 30 minutes per day for a minimum of four days. During this period, both silicon probes and tetrodes were gradually lowered toward the target cell layers. For silicon probe implants, probes were positioned to target the CA1 region, and once the appropriate depth was reached, they were left in place for the remaining days of the experiment. For tetrode implants, tetrodes were adjusted every day to target the CA1 and CA3 pyramidal cell layers. Electrophysiological profiles, including local field potential characteristics such as sharp-wave ripples and gamma oscillations, were used to guide daily placement. On each recording day, tetrodes were lowered into the target layers in the morning to capture ensemble spiking activity and left in place for approximately 1.5–2 hours before recordings began that day. At the end of the recording day, tetrodes were raised by approximately 150 μm to prevent mechanical damage to the hippocampal layers. The following morning, tetrodes were re-adjusted to locate new cells, minimizing the likelihood of recording from the same neurons across days.

Each recording day began with a baseline sleep/rest session (pre-exploration sleep/rest), followed by an exploration session to finish with a post-exploration sleep/rest session (post-exploration sleep/rest). The environments used in these recordings were either open-field arenas for exploration sessions (e.g., 41 cm diameter cylinder, 41 × 41 cm square box; all with 30 cm high walls) or the sleep box (12 × 12 × 28 cm; containing sawdust bedding and nesting material). After placing the mouse in the sleep box, experimenters monitored the animal’s movements and real-time raw electrophysiological signals to confirm that the mouse had started to sleep/rest. In the absence of electromyographic or any other signals to categorize sleep stages, we refer to these offline periods of extended immobility as sleep/rest. Exploration sessions lasted ∼30 minutes; sleep/rest sessions lasted ∼60–90 minutes. All experiments were conducted under dim light conditions (∼20 lux) with low-level background white noise (∼50 dB). In total, the silicon probe dataset included 38 sleep sessions [mean duration (IQR): 82.4 (59.5 – 105.9) minutes per sleep/rest] from 5 mice; the tetrode dataset included 244 sleep sessions [mean duration (IQR): 68.8 (48.0 – 90.2) minutes per sleep/rest] from 13 mice.

### Acquisition of multichannel data and tracking of animal position

Extracellular signals were recorded using an integrated circuit mounted on the animal’s head (model RHD2164, Intan Technologies; http://intantech.com/products_RHD2000.html), which provided a frequency response from 0.09 Hz to 7.60 kHz during the amplification stage. The amplified and filtered signals were digitised at a sampling rate of 20 kHz. These digitised signals were stored alongside additional data streams, including digital pulses indicating the animal’s position (via transistor-transistor logic) and signals from a three-axis accelerometer integrated into the head-mounted device, which measured head movements and provided an additional measure of the animal’s movement. Positional data were obtained using an overhead colour camera (https://github.com/kevin-allen/positrack/wiki), which tracked the movement of LED clusters in three distinct colours affixed to the electrode assembly. These positional signals were captured at a rate of 39 frames per second.

### Processing of Local Field Potential (LFP) signals

LFP signals underwent initial filtering using an 8th-order Chebyshev type I anti-aliasing filter, applied to the wide-band signals sampled at 20 kHz. These filtered signals were then down-sampled to a rate of 1,250 Hz, employing the decimate function within Scipy’s signal submodule (version 1.11.2).

### Sharp-wave/ripple (SWR) event detection

LFPs were first referenced to a channel without CA1 ripples. This differential signal underwent dual stage-filtering: through a ripple-specific bandpass filter (80-250Hz, 4th-order Butterworth Filter), and then through a high-frequency bandpass filter (200-500Hz, 4th-order Butterworth Filter). Instantaneous signal characteristics, including envelopes and phases, were derived using the Hilbert transform.

SWR events were identified by detecting envelope peaks within the ripple band that exceeded a threshold of five times the median value. In instances of multiple peaks within a 20-ms window, only the peak with the highest amplitude was considered. For each event we then identified its onset and offset points - relied on the envelope’s decrease to below half of the established threshold. Analysis extended to quantifying the ripple cycle count within each event by examining phase shifts, with cycle calculation based on the unwrapped phase difference between event onset and offset as previously described ^51^. The mean frequency of each event was calculated by dividing the total cycle count by the event’s duration. Finally, we validated candidate SWR events using four criteria: (1) Ripple band power in the detection channel, calculated as the squared mean amplitude, needed to be double that of the reference channel, ensuring detected events were prominent; (2) The mean frequency of detected events must exceed 80 Hz; (3) Each event should contain at least four complete ripple cycles; (4) Ripple band power should be at least double that of the control high-frequency band. All detected ripples were included in the analyses unless stated otherwise, as appropriate [mean number of detected ripples (IQR): 3,341.6 (2,130.3 – 4,413.5) per sleep/rest]. Namely, for specific analyses requiring preventing influences by overlapping events (which would otherwise contaminate average signals), only isolated ripples were used (i.e., ripple events with no other detected ripple occurring within ±250 ms). This applies to ripple-triggered average signals (e.g., the triggered-average CSD in Figures 1C and S3A-E; the peri-ripple cells’ response in Figure 2B). All the ripple-triggered average signals were referenced to the ripple-envelope peak unless stated otherwise.

### Determination of CA1 pyramidal layer reference channel

To identify the optimal reference channel for SWR events and theta oscillations within the CA1 pyramidal layer, we computed a ripple band score for each channel. This score was calculated by dividing the power in the ripple band (80–250Hz) by the power in an adjacent frequency range (70– 300Hz), using a Welch spectrum (4-second Hann windows overlapping by 50%). The channel with highest score was set as a reference channel within the CA1 pyramidal layer.

### Extraction of theta oscillations from LFPs

To isolate theta oscillations from the LFP data in exploration sessions, we employed the masked Empirical Mode Decomposition method ^71,72^ as implemented by Quinn et al ^73^. With this, we adopted the mask sift procedure using specific mask frequencies set at 350, 200, 70, 40, 30, and 7 Hz, following the parameters optimized in ^74^ grounded in ^75^. For each mask, the amplitude was set to three times the standard deviation of the input signal. This procedure decomposes each LFP signal into oscillatory components termed intrinsic mode functions (IMFs) from faster to lower frequency components. Upon completion of this procedure with the parameters mentioned, six IMFs and a residue were computed, with IMF-6 effectively isolating theta oscillations.

To delineate individual theta cycles, we began by pinpointing peaks and troughs (i.e., the local maxima and minima, respectively) of the obtained theta IMF. The residue of the LFP not captured by the six IMFs was defined as the lower frequency component of the signal and its envelope was used as the amplitude threshold for retaining peaks and troughs for the next step. We then defined each peak-trough-peak sequence as a candidate theta cycle. We took as valid cycles sequences having their peak-trough and trough-peak intervals falling within the 31 to 100 ms range (corresponding to the half period of cycles with frequencies ranging from ∼16 to 4 Hz); and peak-to-peak distance was between 71 ms (equivalent to ∼14 Hz) and 200 ms (equivalent to 5 Hz).

For each validated cycle, we found six control points: the zero-crossing prior to the first peak, the peak itself, the subsequent zero-crossing post the first peak, the trough, and the zero-crossing following the trough. Then, we computed the instantaneous theta phase for each timestamp through a linear interpolation of the control points ^56,76^.

### Wavelet Spectrograms

The spectrograms shown in Figure S2H were generated using the complex Morlet Wavelet Transform. For this analysis, 50 logarithmically spaced frequencies were selected, spanning from 2 Hz to 300 Hz unless otherwise specified. Each wavelet kernel was L1-normalised, meaning the sum of the absolute values of the elements in the kernel was set to 1. This normalisation ensured that the wavelet preserved the relative amplitudes of individual frequency components without amplifying or attenuating them.

### Current source density analysis

We applied current source density (CSD) analysis ^77,78^ to event-triggered LFP signals obtained using linear silicon probes. For ripple-related analyses, we centred LFP signals around the ripple-band envelope peak. Upon configuring these event-based signals, we calculated the current source density at a specific channel *n* and time point using the formula:

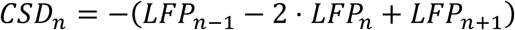

where *n* − 1 and *n* + 1 represent the channels directly above and below channel *n*, respectively. This way, for each ripple event we obtained its CSD signals. In this study, we focused on CA1 channels. We defined the location of *oriens*, *radiatum and lacunosum-moleculare* layers according to the ripple and sharp-wave laminar profiles and electrode spacing, as previously described ^51^. To ensure uniform spatial resolution of CSD measurements across silicon probes with different channel spacing (Table S1), we applied Gaussian kernel smoothing with a standard deviation parameter set to 50 *μm*. For the ripple-triggered average CSD signals in Figures 1C and S3A-E, we only included isolated ripples (see section ‘Sharp wave-ripple (SWR) event detection’).

### Single-ripple CSD signatures

To obtain the CSD signature of each individual ripple, for each channel we computed the mean CSD signal within a 50-ms window around the ripple envelope peak. This way, for each ripple event we obtained a curve which described the average CSD signals during that ripple. We defined this curve as the CSD signature of this ripple event.

### Principal component analysis of single-ripple CSD signatures

To analyse the variance across CSD signature profiles, we applied principal component analysis (PCA). For each recording session, we computed the variance explained by each extracted principal component (PC) as:

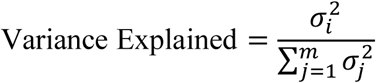

where *σ*_*i*_ is the singular value associated of the *i*^*t*ℎ^ PC and *m* is the total number of PCs extracted. To ensure consistency in the sign of PCs across different recording sessions, we adjusted the sign of the first PC so that it consistently exhibited negative weights within the *stratum radiatum*.

### Identification of Rad^sink^ and LM^sink^ profiles from single-ripple CSD signatures

To investigate the variations in CSD profiles modulated by the first PC across different ripple events, we categorized the distribution of PC1 strengths into three groups, focusing on the extremes of the distribution to capture the most pronounced CSD signatures. Specifically, since PC1 was adjusted to exhibit negative weights in the *stratum radiatum* (indicating that a higher PC1 value corresponds to a stronger current sink in this layer), the ripples with PC1 strengths surpassing the 70th percentile were classified as Rad^sink^ events. Conversely, ripples with PC1 strengths falling below the 30^th^ percentile were designated as LM^sink^ events. The remaining ripples were placed in an ‘baseline’ event category (e.g., used in Figure 1G,I).

### Explained variance of ripple LFPs by laminar CSD

In Figure 1F, we quantified the extent to which the ripple-by-ripple variations in the LFP waveforms from a given recording site placed in the CA1 *stratum pyramidale* can be accounted for by their underlying current source density (CSD) across different CA1 layers, from *stratum oriens* to *stratum lacunosum-moleculare*. To this end, we trained linear decoders to predict for each individual ripple the strength and the sign of the CSD marking each CA1 layer from the LFP recorded in the *stratum pyramidale*. Each ripple-nested *stratum pyramidale* LFP trace was low-pass filtered through a Butterworth filter (4th order, with a cut-off frequency of 30 Hz) to focus specifically on the low-frequency component reflecting the ‘sharp-wave’, which we hypothesized contained all the information to distinguish the underlying current profiles.

Then, the filtered LFP waveforms were standardized through z-score transformation to ensure uniformity in variance and mean across the signals of all ripples. We then applied principal component analysis to reduce the dimensionality of the 200-ms LFP traces down to four principal components, which explained more than 80% of the variance of all ripple-LFP waveforms (Figure S3K). Similarly, the CSD signals for each CA1 layer (*strata oriens, pyramidale, radiatum,* and *lacunosum-moleculare*) were normalized by dividing each CSD signal by its standard deviation calculated over multiple events. This normalization allowed maintaining the polarity information of the CSD signals while ensuring comparability across different magnitudes.

Subsequently, we employed linear regression models to predict the normalized CSD in each CA1 layer from the dimensionality-reduced LFP signals during ripple events. To validate the robustness and generalizability of our models, we performed a cross-validation 20 times (80% training, 20% testing) on the LFP and CSD data. For each iteration, the model was fitted to the training set and then evaluated on the testing set, thereby obtaining a coefficient of determination *R*^2^for each run. The coefficient of determination, *R*^2^, was calculated as:

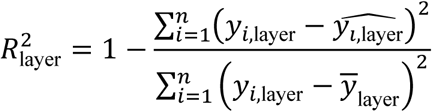

where *y*_*i*,layer_ and 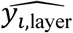 respectively represent the actual and the predicted CSD value for the *i* -th ripple in the specified ‘layer’; *ȳ*_layer_ is the mean of the actual CSD in the specified ‘layer’; *n* is the total number of ripples in each recording session. The variance explained by each model was then determined by averaging the *R*^2^values across all 20 cross-validation iterations, providing a measure of how well the CSD can be predicted from the LFP signals across the different CA1 layers. This cross-validated approach allowed us to assess the predictive power and reliability of our linear decoders in explaining the variance in CSD strength and sign from the LFP recordings during ripple events. The chance level of explained variance of the LFPs was determined by shuffling 500 times the true CSD values and then computing the variance explained by the shuffled data.

### Structure index of LPF waveforms

Using the structure index measure ^79^, we further validated (in Figure S3F) the observed relationship between the ripple-triggered LFP signals recorded in the pyramidal cell layer and the CSD signals obtained in each CA1 layers (in Figure 1F). For each sleep/rest session, we first applied the same pre-processing steps to the LFP and CSD signals described above (see ‘Explained variance of ripple LFPs by laminar CSD’). Then, we computed the structure index for each layer to quantify the interplay between the time profiles of the pyramidal LFPs and the CA1 layer-selective CSD magnitude during ripples. To evaluate the significance of our results, chance levels were determined by shuffling the data 500 times as previously described in the aforementioned study ^79^.

### Single-ripple CSD profile prediction from pyramidal LFP waveform

We applied Linear Discriminant Analysis (LDA) to predict both the polarity (sign) and the magnitude of the *lacunosum-moleculare* (LM) current for each ripple event based on the corresponding *stratum pyramidale* LFP time course. Ripple LFPs waveforms from the pyramidal layer were extracted and pre-processed as explained in section ‘Explained variance of ripple LFPs by laminar CSD’ (n = 117,658 isolated ripples from five mice). To classify each ripple event into the Rad^sink^ versus LM^sink^ category, the corresponding LM CSD was computed within a 50-ms window centred on the ripple peak. This CSD was averaged over three adjacent probe channels: one in *lacunosum-moleculare* and the channels immediately above and below it. To ensure comparability across sleep sessions while retaining information about current polarity, the ripple CSD was normalised by the standard deviation of all ripple events. Ripple events with a normalised negative CSD in the lowest 30th percentile were classified as LM^sink^ ripples, while those in the top 30th percentile were classified as Rad^sink^ ripples. Ripples with CSD values within the 30–70th percentile range were classified as ‘baseline’ ripples (Figure S3H-J), characterised by relatively small current magnitudes.

To assess the robustness of the LDA model, we employed a leave-one-out cross-validation procedure, where each mouse was excluded from the training set in turn, and the model’s accuracy was evaluated on the left-out mouse. To address class imbalance in the training data, we downsampled each class to match the size of the smallest class, repeating this balancing procedure 1,000 times. For each permutation, we also trained a null model with shuffled training labels, disrupting the relationship between pyramidal LFP waveforms and LM currents to estimate chance-level accuracy. In each iteration, we computed the accuracy of both the true model and the null model on the testing data for the left-out mouse. This process provided a chance-normalised accuracy for each left-out mouse, calculated by comparing the mean accuracy of the true models to that of the null models (Figure 1H).

When this model was applied to the tetrode dataset, the pyramidal layer tetrode used for ripple classification was picked as the one with average LFP waveform most similar to the average waveform from the model’s training set. Applying this model to the tetrode dataset resulted in a total of 259,273 Rad^sink^ and 211,505 LM^sink^ ripples from 13 mice [mean number of ripples per sleep/rest (IQR): Rad^sink^, 1,062 (677.8 – 1,417.0); LM^sink^, 866.8 (490.8 – 1,146.3)].

### Spike detection and unit isolation

Spike sorting and unit isolation used an automated clustering approach, leveraging Kilosort (https://github.com/cortex-lab/KiloSort) within the SpikeForest framework (https://github.com/flatironinstitute/spikeforest), as outlined in Pachitariu et al. (2016) ^80^ and Magland et al. (2020) ^81^. For data acquired using tetrodes, KiloSort’s algorithm was adapted to limit templates to channels associated with a specific tetrode bundle and to exclude all other recording channels. Data from all sessions recorded within a single day were concatenated and processed collectively, enabling continuous cell tracking across the day. The clusters generated were manually confirmed by examining cross-channel spike waveforms, auto-correlation histograms, and cross-correlation histograms.

Units selected for analysis consistently exhibited stable spike waveforms, a distinct refractory period in their auto-correlation histograms, and no refractory periods in cross-correlation histograms with other units throughout the day.

### Principal cell versus interneuron classification

Hippocampal principal cells and interneurons were distinguished using features of their spike waveforms, as described previously ^51^. Briefly, waveform consistency for each unit was evaluated using the waveform with the maximum amplitude across tetrode channels for each cluster. To quantify the prominence of a unit’s mean waveform amplitude relative to its spike-to-spike variability, we computed a waveform score:

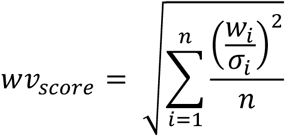

where *w*_*i*_ is the value of the mean waveform at sample *i*, *σ*_*i*_ is the standard deviation at sample *i* across all spikes, and *n* is the number of waveform samples. This metric reflects the relative magnitude of the mean waveform amplitude compared to variability across spikes. Units with a waveform score above 0.75 and fewer than 2% refractory period violations (intervals < 2 ms in the inter-spike interval distribution) were included for further analysis. Putative interneurons and principal cells were then classified based on the trough-to-peak latency of their waveforms. In a prior dataset of 4,000 neurons, the trough-to-peak latency exhibited a bimodal distribution. A one-dimensional, two-component gaussian mixture model was fitted to this distribution, and the intersection of the two components was used as the classification threshold: units with latencies above the threshold were classified as putative principal cells, and those below as putative interneurons. We applied the same inclusion criteria to the principal cells and interneurons in the tetrode dataset of this study. In total, this study includes 2,196 CA1 principal cells and 1,325 CA3 principal cells, with 408 CA1 interneurons and 333 CA3 interneurons, recorded across 83 days in 13 mice [mean number of cells per sleep/rest session (IQR): CA1 principal cells, 26.8 (15.3 – 38.0); CA3 principal cells, 19.8 (14.0 – 25.0); CA1 interneurons, 5.0 (3.3 – 6.0); CA3 interneurons, 5.0 (3.0 – 6.0)]. For analyses involving principal cell correlations or distances between population vectors across ripple types, we addressed the issue of highly sparse spike trains and imbalanced ripple counts across groups by using sleep sessions with at least 250 ripples from each group (Rad^sink^ or LM^sink^) and five principal cells with an average firing rate of at least 0.25 Hz over the entire recording day. This criterion was met by 208 sleep/rest sessions for CA1 principal cells and 171 sessions for CA3 principal cells, resulting in 1,580 CA1 and 866 CA3 principal cells across 13 mice.

### Peri-event time histograms (PETHs)

In Figures 2B and S4G, we constructed PETHs over 400-ms windows, spanning 200 ms on either side of the envelope peak of isolated ripples (see section ‘Sharp Wave-Ripple (SWR) Event Detection’), with a bin width of 0.8 ms. For each cell group (e.g., principal cells or interneurons in CA1 or CA3), we first computed the raw firing rate responses during Rad^sink^ and LM^sink^ ripples. These raw responses were smoothed using a Gaussian kernel (s.d. = 5 ms) and used to calculate the peak firing rate as the maximum rate within a 50-ms window centred on the ripple peak (Figures S4C,F,G). To visualize responses across all cells (Figures 2B and S4G), the responses were then z-scored relative to their mean and standard deviation (s.d.) during Rad^sink^ ripples and further smoothed with a Gaussian kernel (s.d. = 5 ms).

### Preferred ripple phase and phase coherence

In Figure S4D,E, we computed the coupling of CA1 principal cells to ripple oscillations during a single sleep session for each recording day. We first band-pass filtered the LFPs between 90 and 300 Hz using a 4th-order Butterworth filter and estimated the instantaneous phase of the ripple signal during each ripple event, from onset to offset. For each CA1 principal cell, we then calculated the probability of spiking relative to the local ripple phase, using the phase signal from the tetrode where the cell was recorded. The phase range was divided into 24 equally spaced bins between 0 and 2*π*, and the spike-phase probabilities were computed separately for Rad^sink^ and LM^sink^ ripples. From these spike-phase probabilities, we calculated the preferred ripple phase as the angular component of the mean resultant vector of the phase distribution, while the mean phase coherence was derived from the magnitude of this vector, reflecting the strength of phase locking to the ripple oscillations.

### Response similarity between Rad^sink^ and LM^sink^ ripples

For each CA1 and CA3 principal cell, we evaluated the similarity of its activity between Rad^sink^ and LM^sink^ ripples by calculating the Pearson correlation between the firing rates of that cell, or the likelihood of firing at least one spike during ripple events, across the two ripple types (Figures 2C-D and S4I-K). A high correlation coefficient indicated that the cell responded similarly to both ripple types. For example, a cell with a high firing rate (or high spiking probability) in Rad^sink^ ripples showed similar behaviour in LM^sink^ ripples. To compare response similarity between CA1 and CA3 (Figures 2D and S4L), we accounted for differences in cell numbers between the two regions by performing 1,000 permutations. In each permutation, we randomly sampled five principal cells from each region and calculated the response similarity score.

For each sleep session, the average of these permutations provided an estimate of response similarity in both CA1 and CA3. This analysis used sleep sessions with at least five principal cells in the CA region (n = 208 CA1 and 171 CA3 sleep sessions; Figures 2D and S4K).

### Interneurons to Principal cells firing ratio during Rad^sink^ and LM^sink^ ripples

In Figure S4I, we estimated the ratio of interneurons firing rate over principal cells firing rate during Rad^sink^ and LM^sink^ ripples. For both CA1 and CA3, we calculated the mean ripple firing rate of individual interneurons (*rate*_*interneurons*_) and divided it by the mean ripple firing rate of all principal cells (*rate*_*principals*_) recorded on the same day:

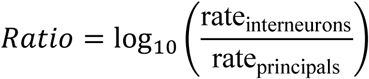

Positive values of this ratio indicated a relatively higher firing rate of interneurons compared to principal cells. This analysis was performed separately for Rad^sink^ and LM^sink^ ripples, and the resulting ratios were compared.

### Population activity discrimination of Rad^sink^ versus LM^sink^ ripples

To determine whether the overall structure of principal cell population activity significantly differed between Rad^sink^ and LM^sink^ ripples, we trained a logistic regression model to predict ripple identity (i.e., whether an event was a Rad^sink^ or LM^sink^ ripple) based on the z-scored ripple-nested population vectors (PVs) containing spike discharge of CA1 or CA3 principal cells (Figure 2E). To control for the number of predictors across CA1 and CA3, we trained these models using multiples of 5 principal cells (i.e., 5, 10, 15, …, N, where N is the maximum number of cells divisible by 5). For each step in the number of cells, we performed 200 permutations, randomly selecting cells for the model. Each model was cross-validated 20 times (80% training, 20% testing), with accuracy measured as the mutual information between the true ripple classes and model predictions (0 bit = chance; 1 bit = perfect prediction). To account for class imbalance, we matched the number of events in each ripple class by resampling the larger class to match the size of the smaller class in each permutation. Additionally, for each step in the number of cells, we trained another model using surrogate PVs, where the coactivity of principal cells was shuffled while preserving the overall firing rate of each cell and the population rate within each ripple (see method section “Spikes shuffling control”).

For each sleep session, we calculated the mean accuracy (or mutual information) across the 200 permutations as a function of the number of cells used for training. This was repeated for the shuffle control models.

These analyses were performed independently for CA1 and CA3. In Figure 2E, we compared the accuracy of the models trained with 15 principal cells, resulting in 129 sleep sessions for CA1 and 89 for CA3.

In Figure S4L-N, we performed an alternative analysis where, instead of using the ripple-nested population vectors, we classified ripple identity based on the entire temporal pattern of spiking activity surrounding each ripple. In each cross-validaiton iteration, we extracted the spike train of each cell within a ±200 ms window around the ripple peak and smoothed this temporal pattern using a Gaussian kernel with a standard deviation of 3 ms. This resulted in a matrix for each cell with dimensions (*N*_*ripples*_ *x N*_*bins*_), where *N*_*ripples*_ is the number of ripples and *N*_*bins*_) is the number of time bins in the ± 200 ms window.

Principal component analysis (PCA) was then applied to this matrix to extract the first two principal components (PCs), which captured the main temporal patterns of the cell across ripples. For each cell, this produced a matrix of dimensions (*N*_*ripples*_ *x* 2) (representing the strength of the first two PCs across all ripples). These matrices were concatenated across all cells to create a larger matrix with dimensions (*N*_*ripples*_ *x* (2 · *N*_*cells*_)), where *N*_*cells*_ is the total number of CA1 (or CA3) principal cells in that recording day. Each PC was then z-scored, and the resulting matrix was used as input for the classifier described above. This procedure was repeated across all 20 cross-validation iterations, ensuring that PCA was independently validated alongside model training. A schematic of this method is shown in Figure S4L.

In Figure S4M, we report the accuracy of this temporal spike pattern classification as a function of the number of CA1 or CA3 cells used for training, and in Figure S4N, we compared the accuracy of CA3 and CA1 models when using 15 cells as predictors. For this analysis, to avoid contamination of the time course waveforms we used only isolated ripples (see section ‘Sharp wave-ripple (SWR) event detection’).

### Spikes shuffling control

We developed a shuffling procedure on spike matrices that preserves both the overall firing rates of individual neurons and the total population activity across time bins while disrupting neuronal coactivity. The original matrix, where each row represents a neuron and each column represents a time bin, was shuffled to maintain the sum of spikes for each neuron (i.e., individual neuron firing rates) and the total number of spikes fired by the population (i.e., overall population firing rate) in each time bin (e.g., a given ripple). This procedure begins by calculating the total spike count for each neuron across all time bins, generating a list of indices corresponding to the times of these spikes. This list is then randomly shuffled to ensure a uniform distribution of spikes across the matrix. The shuffled spike indices are iteratively reassigned to a new matrix, preserving the total number of spikes per neuron (row) and per time bin (column) consistent with the original matrix. This approach controls for individual neuron firing rates and the overall population activity while disrupting the coactivity patterns of neurons within each time bin. This shuffling control is used in Figures 2E, 3H and 4D.

### Neuronal coactivity graphs

To analyse the coactivity of CA1 (or CA3) principal cells during ripples, we constructed the corresponding neuronal graphs. For a given sleep session, we first computed the spike count of each principal cell within a 50-ms window centred at the time of each ripple envelope peak. We then compiled these ripple-triggered activity profiles into a matrix with dimensions given by the number of cells and ripple events (*N*_*cells*_, *N*_*ripples*_). We constructed these matrices independently for ripples classified as Rad^sink^ and LM^sink^. Using these matrices, we constructed coactivity graphs by comparing the ripple-event rates between all pairs of principal cells. To further control for the shared influence of the general network activity between pairs of neurons (*i*, *j*), we fitted a generalised linear model (GLM), thus obtaining the regression coefficient *β* _*ij*_ (Figure 3A):

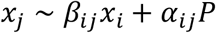

where *x*_*j*_, *x*_*i*_ are the z-scored ripple-nested spike trains of individual neurons *j* (the target) and *i* (the predictor), and *P* is the summed activity of the other *N* − 2 neurons,

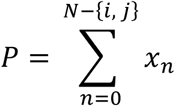

with *α*_*ij*_ weighting the influence of the population contribution to the activity of target neuron *j.* Hence, the recorded neurons (and their coactivity) defined the nodes (and their edges) in the coactivity graphs of each sleep session. We characterized each neuronal graph through its adjacency matrix, *A*, defined as an *N*_*cells*_ × *N*_*cells*_ square matrix that encapsulated the ripple-associated pairwise coactivity interactions across the network, resulting in a (signed and weighted) graph with no self-connections:

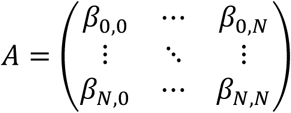

with *β*_*i*,*i*_ = 0 ∀*i in N*, and the additional requirement of symmetric connections 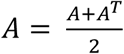 forming an undirected graph.

### Single-neuron coactivity strength

In Figure 3B and S4O-P, we defined the single-neuron coactivity strength as the average pairwise activity correlation of a given node with the other nodes in the weighted graph (n = 1,580 CA1 and 866 CA3 principal cells). As a reference, the strength in a weighted graph can be compared to the degree in a binary graph, which accounts for the number of the node’s neighbours. Here, the strength *S*_*i*_of a node *i* is the average across all the weights *β*_*ij*_ of the edges projected from that node:

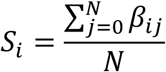

where *N* is the number of neurons *j* that node *i* projects to.

### Structural balance

The combination of positive and negative edges in a network gives rise to both stable and unstable patterns of relationships. In Figures 3E and S4Q, we computed the structural balance of the hippocampal graphs in Rad^sink^ versus LM^sink^ ripples to assess their stability and the coherence of the underlying neuronal coactivity patterns. In this network analysis, triads of neurons are classified as balanced or unbalanced (Figure 3D)^32^. An intuition about such triadic relationships, generalising the notion of clustering coefficient to the signed, weighted setting, arises from social networks where three nodes form a balanced motif either by having three positive edges (e.g., “the friend of my friend is my friend”), or by having two negative ones (e.g., “the enemy of my friend is my enemy”) ^82^. Any triad with an odd number of negative relations would make the motif unbalanced, as different paths between the same pair of nodes would lead to interfering signals. Then, we classified all triads in Rad^sink^ versus LM^sink^ coactivity graphs as balanced or unbalanced based on the signs of their three edges. The structural balance of a graph was defined as the proportion of its balanced triads (Figures 3E and S4Q; n = 208 sleep sessions (graphs) for CA1 and 171 sleep sessions (graphs) for CA3).

### Population-level sparsity

The sparsity (S) of a population firing vector (x) was computed using the Gini index ^83–85^ as:

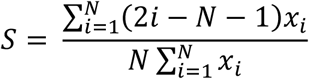

where *x* represents the population vector of spike counts for each principal cell, arranged in ascending order, within a 50-ms time window centred at the ripple peak (either Rad^sink^ or LM^sink^), *N* is the number of simultaneously recorded principal cells (i.e., the length of the vector), and *i* denotes the rank of spike count in ascending order. Population vectors where spike counts are more evenly distributed across neurons have a lower Gini index (indicating lower sparsity), while those where spike counts are concentrated in a few neurons have a higher Gini index (indicating higher sparsity). In Figure 3G, we reported the sparsity for CA1 principal cells (n = 208 sleep/rest sessions).

### Population-level dimensionality

We quantified the intrinsic dimensionality of CA1 ripple-nested activity (i.e., within a 50-ms time window centred at the ripple peak) in Rad^sink^ versus LM^sink^ ripples using the angle based intrinsic dimensionality (ABID) measure ^33^. Notably, ABID is non-linear, making it better suited for capturing complex neural activity patterns.

Briefly, ABID estimates dimensionality (*D*) by analysing the cosine similarity between each ripple population vector (PV) and its k-nearest neighbours (k = 50). For a given sleep/rest session with *M* ripples (calculated separately for Rad^sink^ and LM^sink^), the ripple-nested PVs were z-scored, and the dimensionality of each ripple *m* was computed as:

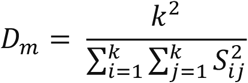

where *S*_*ij*_ is the cosine similarity between the normalised k-nearest PVs of ripple *m*. Dimensionality is inversely related to the concentration of these similarities: when the cosine similarities (*S*_*ij*_) are high, indicating tightly clustered neighbours in the high-dimensional space, the sum of squared similarities 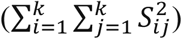 increases, resulting in lower dimensionality. Conversely, when similarities are lower, reflecting a more spread-out distribution of neighbours, the dimensionality is higher. The intrinsic dimensionality for each sleep/rest session was then computed as the average across all *M* ripples. To control for class imbalance between the two ripple types and across sleep/rest sessions, we randomly selected 100 PVs for each ripple type and calculated the intrinsic dimensionality of this subsampled data. This procedure was repeated 1,000 times, and for each sleep session, the mean intrinsic dimensionality for Rad^sink^ and LM^sink^ ripples was defined as the average across these 1,000 permutations (Figure 3I; n = 208 sleep/rest sessions).

### Cross-population vector inclusion of activity motifs

To quantify the degree of inclusion between the coactivity patterns in LM^sink^ versus those in Rad^sink^ ripples, we computed the overlap among their population vectors (PVs). Each ripple-nested PV contained the instantaneous firing activity of CA1 principal cells within a 50-ms window centred around the ripple peak. Each PV was then described as a binary vector where the presence of any spikes from a neuron is denoted by 1, and absence of any spike from a neuron is denoted by 0. This approach allowed constructing an adjacency matrix, where each entry provides a measure of overlap between the activity patterns of different PVs (see Figure S5B-C). Given two population vectors, *PV*_*m*_ = [*p*_*m*1_, *p*_*m*2_, …, *p*_*mn*_] and *PV*_*q*_ = [*p*_*q*1_, *p*_*q*2_, …, *p*_*qn*_], where *n* represents the total number of principal cells in the population, and *p*_*mk*_, *p*_*qk*_ indicate the (binary) firing state of the *k* -th neuron in the pair of ripple-nested population vectors *m* and *q*, respectively. To quantify the overlap in active neurons between two PVs, we computed an asymmetric version of the Jaccard coefficient between *PV*_*m*_ and *PV*_*q*_. Effectively, this metric, denoted as *I*_*m*,*q*_, quantifies the proportion of cells in *PV*_*m*_ that are also active in *PV*_*q*_:

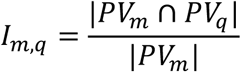

where the ∩ symbol represents the intersection between the two sets of active cells, and the | · | operator represents the cardinality of the set, i.e. the number of (active) cells it contains. Thus, here a value of 1 indicated that all cells active in *PV*_*m*_ were also active in *PV*_*q*_, and zero if none were active in both. Similarly, we defined the proportion of cells in *PV*_*q*_ that are also active in *PV*_*m*_:

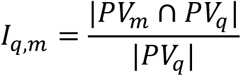

Note the different denominator with respect to the previous equation. It is important to also note that this operation is inherently asymmetric due to the directional nature of the cardinality of the two sets, resulting in *I*_*m*,*q*_ not necessarily being equal to *I*_*q*,*m*_. This asymmetry allowed exploring directional similarities in activity patterns and thus provided insights into the pattern similarities between PVs. To highlight the importance of this directionality (e.g., in Figure 3), we employed notation *I*_*m*,*q*_ = *I*_*m*→*q*_.

### Neural inclusion of LM^sink^ into Rad^sink^ ripples

For each sleep session, we computed the inclusion between each pair (*m*, *q*) of ripple-nested PVs (as presented in the section above). Using pairs of PVs nested in LM^sink^ and Rad^sink^ ripples, we quantified the degree of active cells similarity in the sets of active neurons from LM^sink^ into Rad^sink^ ripples. This provided insights into the directional pattern similarities and differences between these ripple types. To account for the potential bias due to an imbalance in the number of instances between LM^sink^ and Rad^sink^ ripples, we standardized the comparison by matching the number of PVs from each class to the smaller of the two, denoted by *D*. Then, the mean overlap of LM^sink^ into Rad^sink^ PVs (*I*_*LM*→*Rad*_) was calculated as the average across these subsampled *D* PVs.

To further mitigate potential biases arising from subsampling specific sets of PVs, we adopted a resampling strategy. In each iteration, the inclusion *I*_LM→Rad_ was recalculated using a randomly selected subset of *D* PVs from the class with the larger number of instances. This procedure was repeated across 1,000 iterations to account for variability within the data.

Additionally, in each iteration, we generated a chance level by applying a shuffling control that randomized the active cell distribution within LM^sink^ ripples while leaving Rad^sink^ ripples unchanged (see ‘Spikes shuffling control’ section). Specifically, for each LM^sink^ ripple, the active cells were randomized while preserving both the original number of active cells (reflecting the ripple’s sparsity) and the total number of LM^sink^ ripples in which each cell was active. To assess the significance of the observed inclusion, we compared the mean actual inclusion values to those derived from the shuffled data (*I*_*LMs*ℎ*uffled*→Rad_) (Figure 3H; n = 208 sleep/rest sessions). This method enabled us to evaluate whether the overlap *I*_LM→Rad_ resulted from specific cells being active in both LM^sink^ and Rad^sink^ ripples or if it emerged randomly due to the higher sparsity in LM^sink^. It is important to note that we avoided direct comparison between *I*_LM→Rad_ and *I*_*Rad*→*LM*_ because the sparsity differences between the two ripple sets could bias their respective overlaps.

### Classification of CA1 principal cells into deep and superficial

We classified CA1 principal cells as either deep or superficial, following the approach described in previous work ^51^. Briefly, we extracted LFP features from the tetrode where each principal cell was recorded. These features were then projected onto a linearized trajectory, which estimates the cell’s depth within the pyramidal layer specifically, whether the associated tetrode is closer to the *stratum radiatum* or the *stratum oriens.* As illustrated in Figure 4A, the depth distribution of CA1 principal cells along this trajectory is bimodal. Using this distribution, we classified cells as superficial or deep, with a threshold depth value of 6 separating the two groups, resulting in a total of 1,353 deep and 843 superficial cells. For analyses where the criteria described in ‘Principal cell versus interneuron classification’ were applied, the numbers were refined to 1,100 deep and 480 superficial cells.

### Change in firing rate during ripples relative to baseline

In Figures 4C (n = 1,100 deep and 480 superficial cells), we compared the change in firing rate deep and superficial principal cells relative to a pre-ripple baseline. This allowed comparing cells with different firing rates in ripples (i.e., deep versus superficial CA1 principal cells. To compute this measure, we calculated the mean instantaneous firing activity of each cell within a 50-ms window centred around the ripple peak (rate_ripple_). To estimate the baseline firing rate (rate_baseline_), we computed the mean firing activity in a time window from −200 ms to −100 ms relative to the ripple peak. Then, independently for Rad^sink^ and LM^sink^ ripples, we calculated the increase in firing rate as:

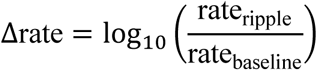

where positive values of Δrate indicate an increase in firing rate relative to the pre-ripple baseline, while negative values indicate a decrease. For this analysis, we included only isolated ripples to avoid contamination of the baseline rate (see section ‘Sharp wave-ripple (SWR) event detection’).

### Additional cells active in Rad^sink^ but not in LM^sink^ ripples

In the analysis presented in Figure 4D, we investigated cells exhibiting preferential firing for Rad^sink^ but not LM^sink^ ripples (see Figure 3). We first identified the cells active in each ripple, as described in section “Cross-population vector inclusion of activity motifs.” For each cell, we then calculated the conditional probability of it being active in a Rad^sink^ ripple given that it was inactive in an LM^sink^ ripple. To control for the intrinsic firing probability of each cell across ripples, we estimated a chance level for each cell using a shuffling procedure. This procedure independently shuffled the coactive neurons during Rad^sink^ and LM^sink^ ripples while maintaining the total number of ripples in which the cell was active and preserving the sparsity of each ripple (see “Spikes shuffling control” section). The shuffling process was repeated 200 times, and the resulting chance levels were used to z-score the observed conditional probabilities (Figure 4D, n = 1,100 deep and 480 superficial cells).

### Offline reactivation

To analyse offline reactivation of CA1 or CA3 (Figure S6A,B) principal cell spiking patterns during sleep/rest, we first trained a linear regression model using awake theta-cycle activity from the exploration session of that recording day. Specifically, the model was trained to predict the firing rate of a target cell based on the activity of four other cells, ensuring that none of the predictor cells were recorded on the same tetrode as the target cell. This criterion allowed avoiding biases in correlations that could arise from shared global tetrode activity. Model accuracy was evaluated as the Pearson correlation between the actual firing rate of the target cell and the firing rate predicted by the model. To ensure robustness, we performed 100 bootstrap iterations, randomly selecting four predictor cells from the pool of cells recorded on the same day for each iteration. Next, we applied these cross-validated models to the z-scored ripple-nested (using a 50-ms window centred around the ripple peak) firing activity of the target cell during pre-exploration and post-exploration sleep/rest sessions. For each cell, the mean accuracy across the 100 cross-validated models was calculated separately for pre-exploration sleep and post-exploration sleep, providing two model accuracy values. Reactivation was defined by comparing the overall accuracy across all cells in post-exploration sleep to the corresponding accuracy in pre-exploration sleep. Specifically, significant reactivation was identified if the post-exploration accuracy was higher than the pre-sleep accuracy (1-tailed paired bootstrap tests; Figure S6A,B; n = 1,580 CA1 and 866 CA3 principal cells). This analysis was performed independently for Rad^sink^ and LM^sink^ ripples during both pre- and post-sleep.

### Deep and superficial cells reactivation

In Figure 4E-G, we computed the offline reactivation of awake spiking patterns for deep and superficial cells. This analysis followed the same approach described in the “Offline reactivation” section, with an additional criterion: predictor cells and the predicted cell were required to belong to the same CA1 pyramidal sublayer. Specifically, a set of four superficial cells was used to predict the activity of another superficial cell, and a set of four deep cells was used to predict the activity of another deep cell, as illustrated in Figure 4E. Together with the existing criterion that predictor cells and the predicted cell must be recorded on different tetrodes (see section “Offline reactivation”), this sublayer-specific restriction reduced the pool of eligible cells to n = 1,097 deep and 478 superficial CA1 principal cells.

### Coactivity pattern stability in LM^sink^ and Rad^sink^ ripples

To measure whether the coactivity patterns nested in Rad^sink^ and LM^sink^ ripples changed from pre-exploration to post-exploration, we first isolated ripple-nested population vectors (PVs) that contained the instantaneous firing activity of CA1 principal cells within a 50-ms window centred around the ripple peak. This was done separately for pre-exploration sleep and post-exploration sleep/rest sessions. Using the z-scored PVs from pre-sleep, we trained a generalized linear model (GLM) to predict the firing response of each CA1 principal cell based on the activity of the rest of the population during each ripple. This process was repeated independently for Rad^sink^ and LM^sink^ ripples, yielding two models for each cell (one for LM^sink^ and one for Rad^sink^ ripples). Each model was cross-validated 20 times (80% training, 20% testing), and accuracy was assessed as the mean correlation between the predicted and true activity of the testing set for each iteration: *r*_*pre*,*i*_. To estimate the chance level for each GLM, we used a shuffling procedure that randomized the cell IDs within each PV 500 times prior to training, breaking cell correlations but preserving the overall rate for each PV. To measure coactivity changes from pre- to post-exploration rest, we applied the cross-validated model of each cell to the (z-scored) PVs from post-sleep and measured its accuracy (*r*_*post*,*i*_). For both Rad^sink^ and LM^sink^ ripples, we quantified the coactivity stability for cell *i* as:

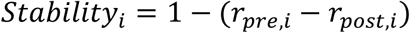

where *r*_*pre*,*i*_ and *r*_*post*,*i*_ are the accuracy values in pre- and post-sleep of the GLM of cell *i* which were normalised with respect to the chance level estimated through the shuffling procedure mentioned above (Figure S6G; n = 140 pre- and post-sleep pairs). A high stability score (∼1) indicates that the model’s accuracy was similar between pre- and post-sleep, reporting that coactivity patterns did not undergo significant cross-session changes. Conversely, a low stability score reflects a substantial change in coactivity from pre- to post-sleep.

### Reactivation of prior and recent coactivity motifs

To determine whether Rad^sink^ and LM^sink^ ripples preferentially reactivated (*i*) the CA1 neural patterns containing the spike discharge from the neurons that were selectively coactive in the most recent wakefulness (i.e., neurons coactive during the open-field exploration that day but not during the pre-exploration sleep/rest before) and/or (*ii*) the prior neural patterns of neuronal coactivity (i.e., the pre-existing patterns, which were already present in the pre-exploration sleep/rest), we first constructed coactivity graphs (as described in section “Neuronal coactivity graphs”). For the pre-sleep coactivity graph, we sampled an equal number of Rad^sink^ and LM^sink^ ripples, establishing a coactivity baseline. Similarly, we constructed the coactivity graph from the awake session, using spiking activity nested in waking theta cycles instead of offline ripples. Thus, for each recording day, we obtained two coactivity graphs: *A*_*pre*_ and *A*_*t*ℎ*eta*_. To isolate coactivity motifs recently expressed during wakefulness but absent in pre-sleep, we modelled the change in coactivity from pre-sleep to wakefulness as a linear transformation:

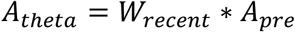

From this, we computed the transformation matrix:

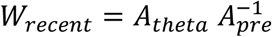

where *W*_*recent*_ represented the coactivity motifs that emerged during wakefulness. Similarly, to identify coactivity motifs present in pre-sleep but absent in the subsequent wakefulness, we used the transformation:

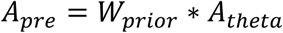

Thus, the transformation matrix is given by:

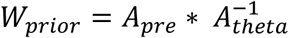

We then evaluated the strength of the recently-expressed coactivity motifs and the prior coactivity motifs in both Rad^sink^ and LM^sink^ ripples of the subsequent post-exploration sleep. For this, we first extracted the post-sleep population PVs that contained the instantaneous firing activity of CA1 principal cells within a 50-ms window centred around the ripple peak. Applying the transformation matrices *W*_*recent*_ and *W*_*prior*_ to the original post-sleep ripple- PVs effectively projected the original coactivity space onto subspaces representing either recent or prior motifs. For each PV, we computed the dot product between the PV in this transformed space and in the original coactivity space, yielding a reactivation score for each ripple. A high reactivation score indicated that the original activity space accurately reflected the transformation (either recent or prior), while a score of zero indicated orthogonality to the transformation space. Reactivation scores were independently computed for Rad^sink^ and LM^sink^ ripples during post-sleep, and the mean within each ripple group was defined as the reactivation score for that ripple type in that sleep session.

### Simulation of prior and recent coactivity motifs

In Figure S6C-F, we validated the method described in the section “Reactivation of prior and recent coactivity motifs” using in silico simulations. For this, we first simulated a neuron ensemble pre-sleep activity matrix containing two pre-configured patterns of coactive neurons (prior motifs). Next, we simulated an awake exploration by incorporating additional patterns of coactive neurons (recent motifs). Specifically, one pre-sleep motif did not change while the second was modified by the integration of an additional neuron, and a third motif was newly formed during the simulation. Using these simulated data, we constructed two sets of post-sleep ripples. The first set of ripples (referred to as “rigid ripples”) expressed only the same neuronal motifs as those seen in pre-sleep. The second set of ripples (referred to as “plastic ripples”) also included these motifs along those observed formed during the awake exploration. We then applied to this “ground truth” *in-silico* data the method described in “Reactivation of prior and recent coactivity motifs” to assess successful extraction of prior and recent motifs from the two post-sleep matrices. We computed the strength of these motifs in each post-sleep scenario to evaluate the accuracy of the method.

### Contribution of deep versus superficial cells to reactivation of coactivity patterns

In Figures 5C-D, we measured the contribution of deep and superficial cells to the reactivation of recently expressed and prior patterns of neuronal coactivation (see “Reactivation of prior and recent coactivity motifs”). Specifically, we computed the post-sleep strength of recent and prior patterns within groups of five deep cells or five superficial. To cross-validate the selected cells, we performed 500 permutations by randomly sampling five deep cells or five superficial cells from the pool of principal cells recorded that day. The mean across these permutations was used to define the reactivation strength for each sublayer in each ripple.

Finally, for each pre- and post-sleep pair, we compared these reactivation values (prior- or recent-motifs) between deep and superficial cells. To ensure a fair comparison and account for potential imbalances in the number of cells (or the absence of cells from a specific sublayer) on a given recording day, we required at least five deep and five superficial principal cells (Figures 5C-D, n = 43 pre- and post-sleep pairs from 6 mice).

### Changes in reactivation over hour-long timescale

In Figures 5E-F and S6H-J, we quantified changes in the reactivation of prior or recent coactivity motifs over hour-long timescale (see section “Reactivation of prior and recent coactivity motifs”). For this analysis, we included only pre- and post-sleep pairs where the post-exploration sleep lasted at least 1 hour (n = 118 pre- post-sleep pairs from 13 mice). For each session, we calculated the mean reactivation of prior and recent motifs during Rad^sink^ or LM^sink^ ripples, dividing the first hour of sleep into 10-minute bins. To ensure comparability of reactivation values across days with different numbers of recorded cells, we normalised the reactivation by subtracting the global mean reactivation for each motif (i.e., prior or recent), irrespective of ripple type (Rad^sink^ or LM^sink^). To assess whether reactivation significantly changed over time, we used an exponential fit model (see “Exponential fit of reactivation over time” section).

### Exponential fit of reactivation over time

To determine whether reactivation changed exponentially over time, we modelled the temporal changes using an exponential decay function. First, we concatenated data from all sleep/rest sessions, creating two vectors: one representing the time of each bin and the other representing the corresponding reactivation values (for prior motifs, recent motifs, Rad^sink^ ripples, or LM^sink^ ripples). Changes in reactivation over time were then modelled as:

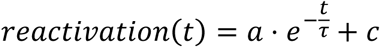

where *a* is the amplitude (scaling factor), *τ* is the time constant (decay parameter), and *c* is the offset or baseline. We used the optimize submodule from the SciPy Python package to fit this function to the concatenated session data. A hyperparameter search was conducted to find the best initial estimates for *a*, *τ*, and *c*, maximising the fit between the model and the data. The goodness of fit was quantified using the Pearson correlation *r* between the observed reactivation values and the model predictions. To estimate the mean fit score (*r*) and time constant (*τ*), we conducted a cross-validation by fitting the exponential function using a bootstrapped subset of 80% of the pre- and post-sleep pairs. This procedure was repeated over 10,000 resamples, generating distributions of estimates for the *r* and *τ*. To assess the significance of the fit, we also computed the fitness score for models with shuffled time information. This shuffling procedure was repeated 10,000 times, and the p-value was calculated as the proportion of shuffles that achieved a fitness score equal to or greater than the actual score. A fit was considered significant if p < 0.05. To avoid potential bias in detecting or fitting trends due to changes in sleep quality at the beginning of the session, we excluded the first 10 minutes of each sleep session from the analysis.

### Reactivation over time controlling for ripple occurrence frequency and ripple population sparsity

In Figure S6H-K, we evaluated whether the observed gradual changes in reactivation over time during LM^sink^ ripples (see Figure 5F) were genuinely time-dependent (i.e., related to the ripple occurrence time in sleep/rest session) or driven rather related to changes in ripple occurrence frequency (i.e., how many ripples per unit of time) or ripple population sparsity (i.e., how many active neurons per ripple) (Figure S6H,J). To address this, we concatenated the time-binned reactivation values of recently-expressed motifs (during Rad^sink^ or LM^sink^ ripples) and trained a linear regression model to predict reactivation at each time bin (see section ‘Changes in reactivation over hour-long timescale’). The predictors included: (1) the ripple occurrence time (i.e., the time of the bin), (2) the ripple occurrence frequency (mean number of ripples per minute in each time-bin), and (3) the ripple population sparsity (calculated as the mean proportion of active cells in ripples in each time-bin). Given the non-linear relationship between time and reactivation (see Figure 5E,F), we applied a log10 transformation to the ripple occurrence times to linearise this relationship. To avoid potential biases introduced by the quality of sleep, we excluded the first 10 minutes from each session. The model was trained and cross-validated 20 times (80% training, 20% testing), with accuracy measured as the Pearson correlation between the observed reactivation values and the model’s predictions. To establish a chance level, we shuffled the reactivation values 25,000 times and compared the model’s accuracy to these shuffled data (Figure S6I).

To determine the contribution of each feature to the model’s accuracy, we trained additional models where individual features were shuffled 25,000 times, thereby removing specific information from the shuffled feature. The gain in accuracy for a given feature was calculated as the difference between the accuracy of the original model and that of the feature-shuffled model (Figure S6J). A high accuracy gain for a feature indicated strong predictive power, showing that the feature contributed significantly to changes in reactivation. The significance of each feature was assessed by calculating p-values as the proportion of shuffled models with a gain less than 0 (indicating the feature significantly contributed to the model).

### Data and statistical analyses

Data analyses were conducted using Python versions 3.6 and 3.10, incorporating the following packages: DABEST ^87^, scikit-learn ^88^, NetworkX ^89^, NumPy^90^, SciPy ^91^, Stats-Models ^92^,Matplotlib ^93^, Pandas ^94^, and Seaborn ^95^. Symmetric distribution assumptions underpinned the two-sided statistical tests, visualised using Gardner-Altman and Cumming plots from the DABEST framework. These plots illustrate effect sizes by comparing mean or median differences across groups. Each plot consists of two panels: the top shows raw data distributions with group means ± SEM (unless stated otherwise), and the bottom shows differences relative to a reference group, calculated from 5,000 bootstrapped samples. Black dots represent the mean, black ticks indicate 95% confidence intervals, and bootstrapped error distribution curves are included. Statistical comparisons included t-tests against a mean (to compare one distribution to a fixed value) and one-way ANOVA for multiple conditions, followed by a Tukey post-hoc test. To compare two conditions, bootstrap tests were employed. These tests, which accommodated both paired and unpaired comparisons, estimated the bootstrapped mean difference (either absolute or as a percentage relative to one of the two variables) by resampling the data 100,000 times (unless stated otherwise) with replacement. For paired comparisons, indices were resampled to preserve the relationship between pairs, whereas for unpaired comparisons, each condition was resampled independently. P-values for these tests were computed numerically, under the null hypothesis of zero difference. For one-sided tests, the p-value was calculated as the proportion of bootstraps where the difference distribution was either greater than or less than zero, depending on the test direction. For two-sided tests, the p-value was determined by multiplying the smaller proportion of bootstraps below or above zero by two. Unless otherwise specified, all p-values from bootstrap tests were two-sided. All confidence intervals (95% CI) were calculated via bootstrapping with 100,000 resamples (unless stated otherwise). For each interval, data were resampled randomly with replacement, and the 2.5^th^ and 97.5^th^ percentiles of the bootstrapped distributions determined the lower and upper bounds of the CI.

## Supplementary Figures and legends

**Figure S1:**
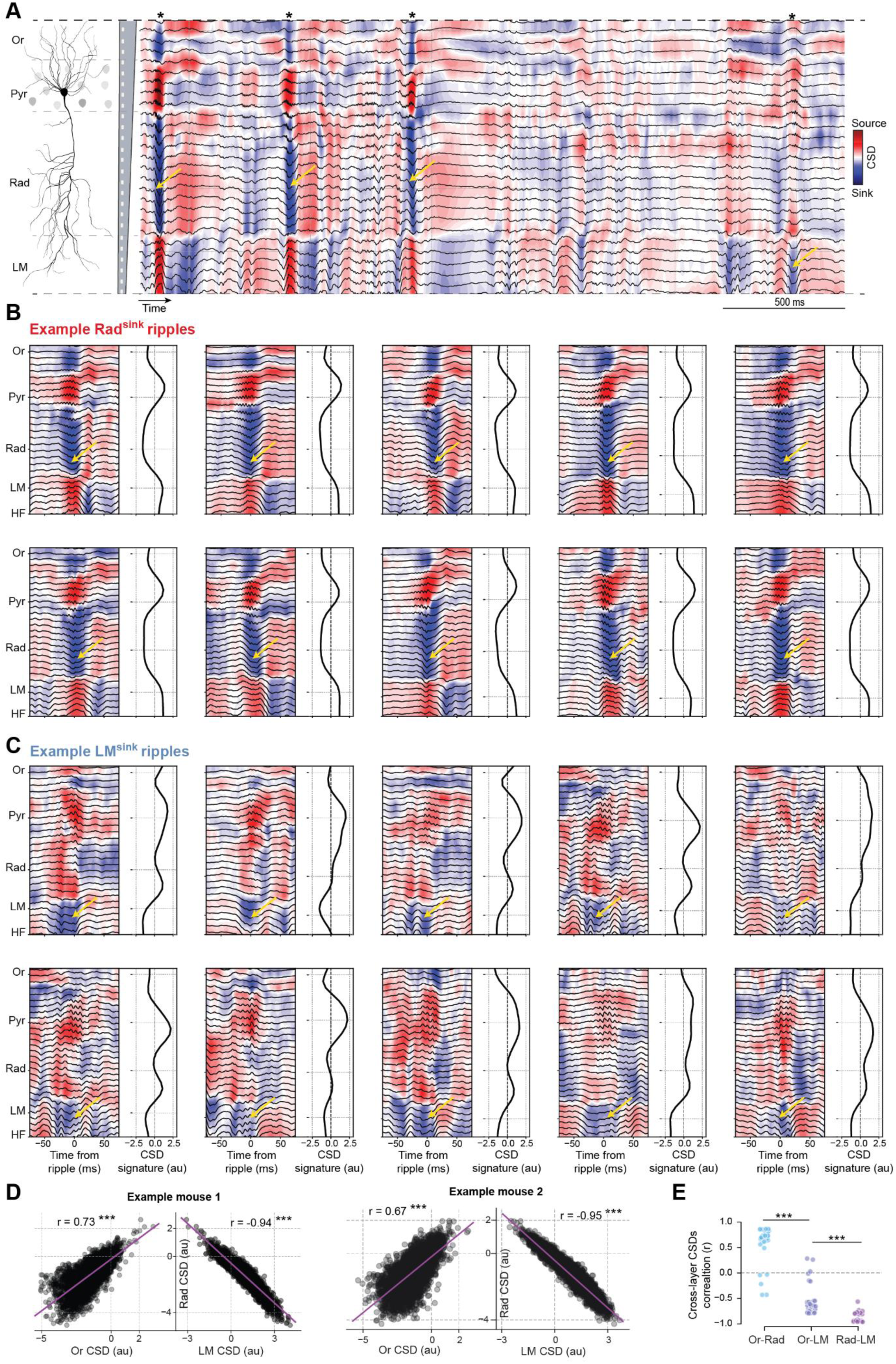
Individual Rad^sink^ and LM^sink^ ripples have distinct CSD profiles. **(A)** Example silicon probe recording (a 3-sec sample for clarity) spanning the somato-dendritic axis of CA1 principal cells to show the instantaneous CSD (coloured hit map; red, source; blue, sink) and local field potential waveform (black traces) across the different neural layers (*strata oriens*, Or; *pyramidale*, Pyr; *radiatum*, Rad; *lacunosum-moleculare*, LM) during a sleep session. Each black asterisk indicates an individual ripple, and each yellow arrow marks a current sink (in *stratum radiatum* or *stratum lacunosum-moleculare*). **(B-C)** Shown are raw CSD signals (*left*, hit maps) with LFP waveforms (*left*, black traces), and the associated CSD signature (*right*, black curve) for ten Rad^sink^ ripples (B) and ten LM^sink^ ripples (C). Yellow arrows highlight a current sink in stratum radiatum or stratum lacunosum-moleculare, as appropriate. **(D)** Relationship between individual ripple-CSD values in *stratum oriens, radiatum, and lacunosum-moleculare*. Each dot represents a ripple, with coordinates indicating the average CSD values across the three layers for an example sleep session. Purple lines represent the best linear fit; numbers report the Pearson correlation coefficients of ripple-CSDs across two layers. **(E)** Distribution of mean Pearson correlation coefficients of ripple-CSDs across different layers. Each dot corresponds to a sleep session (p = 5×10^-48^; one-way ANOVA; all pairwise Ps < 10^-6^, Tukey post-hoc test). *p < 0.05, **p < 0.01, ***p < 0.001.

**Figure S2:**
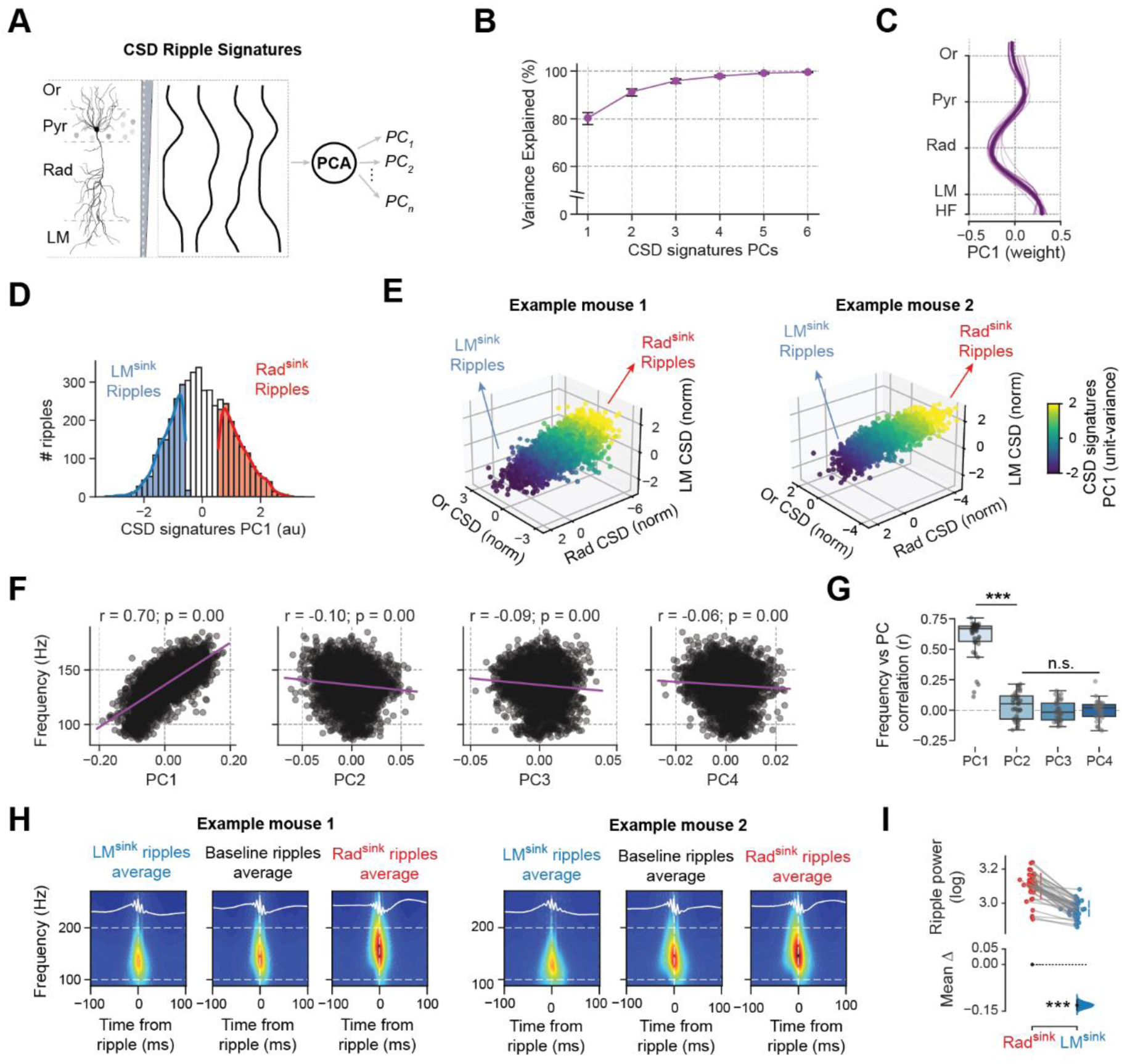
Rad^sink^ and LM^sink^ ripples can be distinguished by Principal Component Analysis of laminar CSD profiles. **(A)** We applied Principal Component Analysis (PCA) on the CSD signatures (spanning the CA1 radial axis) of all individual ripples, as a dimensionality reduction method to identify CSD-signature components that explained the most variance across individual ripple events. **(B)** Mean variance explained by each CSD signature principal component (PC) [mean variance explained of PC1 (95% CI): 80.3 (77.6 –82.6) %]. *Black error bars*, 95% CI. **(C)** Laminar organisation of PC1 weights. Note the larger contributions from sinks in stratum radiatum, which align with the averaged CSD profile reported in Figure 1B. *Thinner lines*, individual sleep/rest sessions. *Thicker line*, mean PC1 across all sessions. **(D)** Shown for the ripples of an example sleep session is the distribution of PC1 strength. Note the significant cross-ripple variability. Ripples at the lower and upper extremes of the distribution (bottom and top 30%) correspond to stronger current sinks in lacunosum moleculare (LM^sink^ ripples, blue) and radiatum (Rad^sink^ ripples, red), respectively. **(E)** 3D scatter plots of ripples from sleep sessions in two example mice, with coordinates representing instantaneous ripple-CSD values in stratum oriens, radiatum, and lacunosum-moleculare. CSD values normalised to be unit-variance. Points are color-coded by PC1 strength, highlighting the continuum of ripples in this lower-dimensional space. Ripples with stronger PC1 values align with the two profiles identified in (D), corresponding to Rad^sink^ or LM^sink^ ripples. **(F)** Scatter plots showing the relationships between ripple frequency and CSD principal component (PC) strength for an example sleep session. Each point represents a ripple. Pearson correlation values computed between ripple frequency and the strength of each PC. Only PC1 showed a strong correlation with ripple frequency. **(G)** Boxplot and scatter plot showing the mean Pearson correlation between ripple frequency and strength of each PC. Each dot represents a sleep session. Note that the stronger PC1 strength, indicative of a stronger sink in the radiatum, is associated with higher ripple frequency (p = 1×10^-57^, one-way ANOVA; PC1 versus PC2: p = 6.9×10^-14^, Tukey post hoc test). **(H)** Spectrograms of CA1 pyramidal LFPs from two example mice across ripple types. For each mouse, the panels (from left to right) show the average spectrogram (with LFP waveform, white trace) for LM^sink^, Baseline, and Rad^sink^ ripples within a ±100 ms window centred on the ripple-envelope peak. Note that from left to right the average ripple power increases as well as the mean ripple frequency. **(I)** Estimation plot showing the effect size for the difference in ripple power, which is significantly higher in Rad^sink^ than LM^sink^ ripples (p < 10^-5^; bootstrap test). Each dot represents a sleep session. *p < 0.05, **p < 0.01, ***p < 0.001.

**Figure S3:**
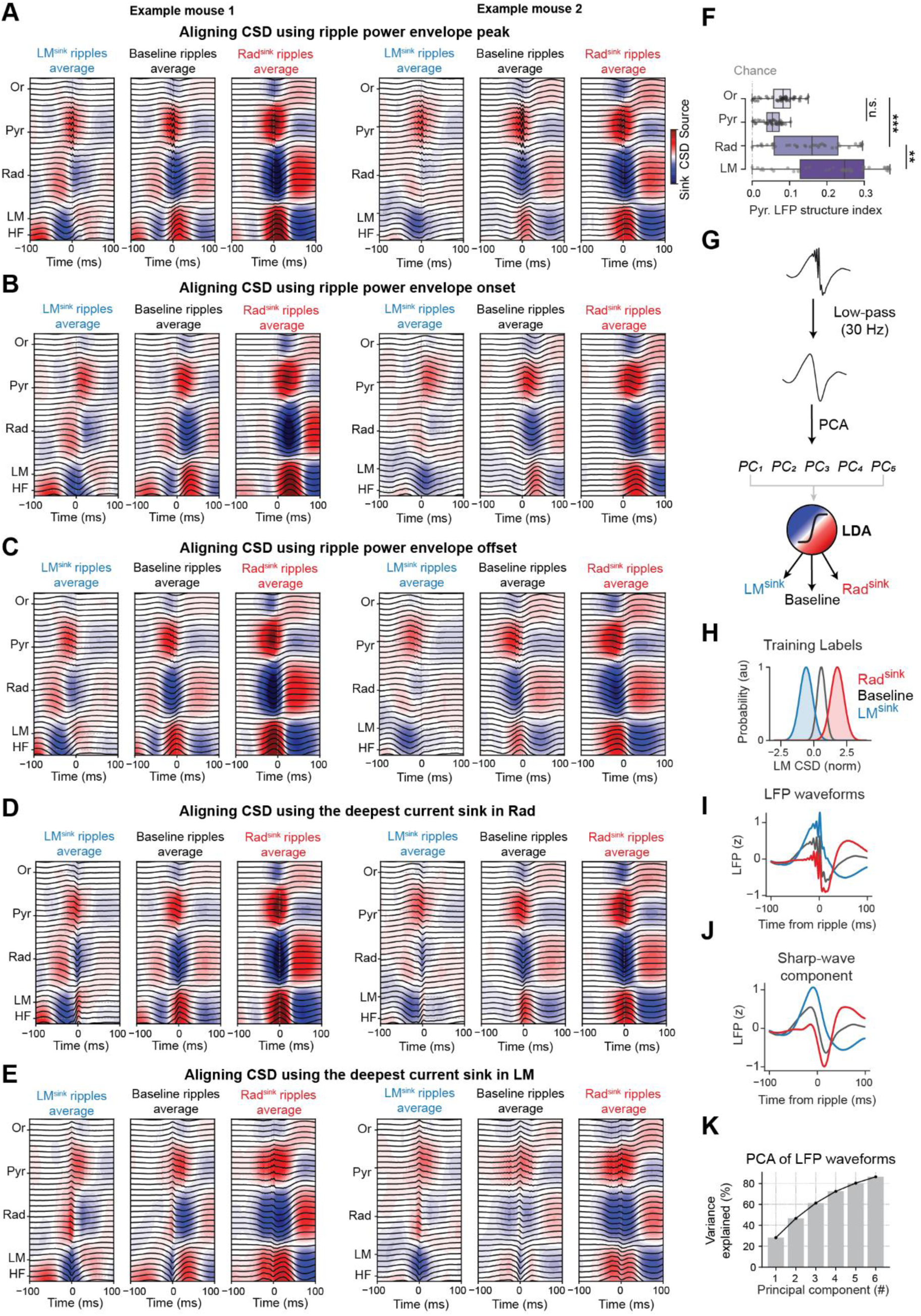
Rad^sink^ and LM^sink^ ripple profiles emerge consistently and are predicted from the pyramidal LFP waveform. (A-E) By aligning CSD with different timestamps within the ripple event, we observed that the Rad^sink^ and LM^sink^ profiles consistently emerged regardless of how the LFP and the CSD signals were referenced. Shown are triggered average CSD and LFP signals across CA1 layers during LM^sink^, Baseline, and Rad^sink^ ripples (see methods) from two example mice (left and right). Note that Rad^sink^ and LM^sink^ ripple profiles were consistent regardless of whether LFP and CSD signals were triggered using the ripple envelope peak (A), onset (B), offset (C), or ripple current sinks in the radiatum (D) or lacunosum-moleculare (E) layers. To avoid contamination of the averages by overlapping events, this analysis only includes ripples that were isolated (i.e., with no other ripple within ± 250 ms). **(F)** Pyramidal LFP Structure Index ^79^. Boxplot showing the structure index (SI) of LFP ripple waveforms in CA1 pyramidal layer explained by the CSD values across CA1 layers. SI normalized by subtracting the average SI by chance (using surrogate models; see Methods). Note that currents in lacunosum-moleculare captured significantly more LFP variance than other layers (p = 2.7×10^-17^, one-way ANOVA; SI in LM CSD versus Rad CSD: p = 2.4×10^-3^, Tukey post hoc). Each dot represents a sleep session. Vertical gray dashed line indicates chance level. **(G-K)** Rad^sink^ and LM^sink^ ripple profiles are predicted from the ripple waveform in *stratum pyramidale* LFPs. The schematic of the linear discriminant analysis (LDA) model used to classify Rad^sink^ and LM^sink^ ripples based on pre-processed LFP waveforms is shown in (G). Ripple events were divided into three classes (Rad^sink^, Baseline, and LM^sink^ ripples) based on digitised current values in the lacunosum-moleculare (H), the layer with the highest predictive power (see Figures 1F and S3F; and Methods). In (H), the red distribution represents Rad^sink^ ripples, the gray distribution represents Baseline ripples, and the blue distribution represents LM^sink^ ripples. For each ripple, the pyramidal LFP waveforms were z-scored (I) and low-pass filtered to isolate their ‘sharp-wave’ component (G,J; see method section ‘Single-ripple CSD profile prediction from pyramidal LFP waveform’). The triggered average LFP waveforms and ‘sharp-wave’ components for the three ripple classes, as defined by the LM CSD values, are shown in (I) and (J). The filtered waveforms were then decomposed into five principal components (PCs), which collectively explained more than 80% of the variance across all ripples (K). These five PCs were used to train the LDA classifier (G) to predict the class of each ripple (i.e., Rad^sink^, Baseline, or LM^sink^). *** p < 0.001.

**Figure S4:**
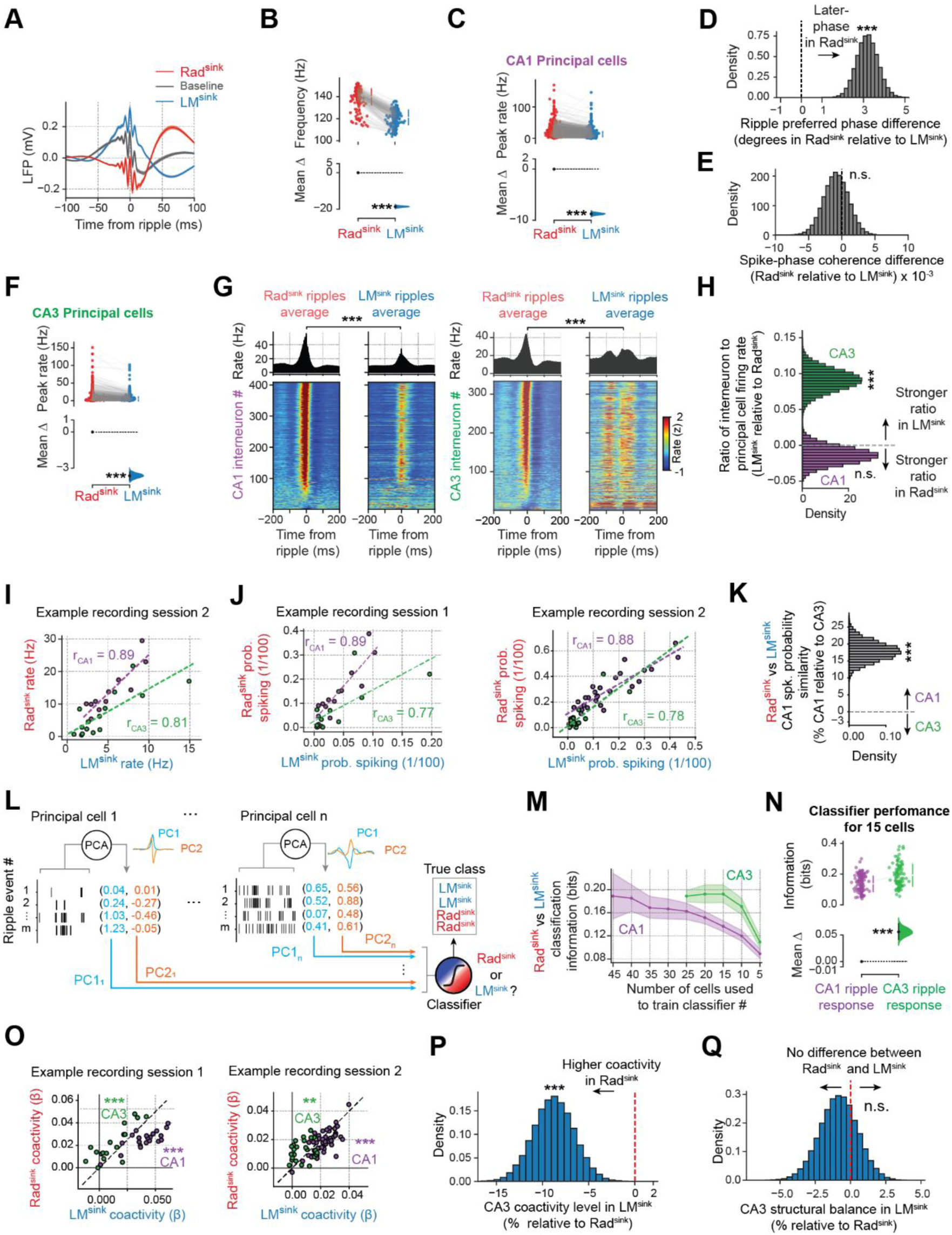
Differential CA1 and CA3 neuron responses in Rad^sink^ versus LM^sink^ ripples. **(A)** Applying the silicon probe-validated CSD ripple-type classifier to the tetrode-recorded dataset retrieved LFP traces (referenced to the highest ripple-peak) for Rad^sink^, baseline, and LM^sink^ ripples that are consistent with those observed in the silicon probe-recorded dataset (Figure 1D). **(B)** Estimation plot showing the effect size for the difference in the ripple frequency of events classified as Rad^sink^ and LM^sink^ by the ripple-type classifier in the tetrode-dataset (see Figure S3G-K). These match the ground-truth differences observed in the silicon probe dataset [Figure 1E; mean frequency (95% CI): Rad^sink^, 139.1 (138.9 – 140.9) Hz; LM^sink^, 122.1 (120.1 – 122.1) Hz; p < 10^-5^; paired bootstrap test]. Each dot is a sleep session. **(C)** Estimation plot showing the effect size for the difference in the peak firing rate of CA1 principal cells during Rad^sink^ versus LM^sink^ ripples. Each point represents a CA1 principal cell. **(D-E)** Mean difference between Rad^sink^ and LM^sink^ ripples (relative to LM^sink^) for ripple preferred firing phase (D) and spike-phase coherence (E) of CA1 principal cells. Overall, CA1 principal cells fired at earlier phases during LM^sink^ ripples (p < 10^-5^; paired bootstrap test), while keeping similar phase coherence across the two ripple types (p = 0.65; paired bootstrap test). **(F)** Same as (C), but for CA3 principal cells. **(G)** Triggered average responses of interneurons in CA1 (left) and CA3 (right). *Top panels*, population average response in Rad^sink^ and LM^sink^ ripples. *Bottom panels*, firing response (z-scored) of individual interneurons. Cells sorted based on their firing rates in Rad^sink^ ripples. CA1 interneurons exhibited stronger responses in Rad^sink^ compared to LM^sink^ ripples (mean peak rate (95% CI): Rad^sink^, 53.04 (48.36 – 57.85) Hz; LM^sink^, 31.57 (28.55 – 34.70) Hz; p < 10^-5^, paired bootstrap test). Similarly, CA3 interneurons showed heightened activity in Rad^sink^ ripples (mean peak rate (95% CI): Rad^sink^, 41.48 (37.02 – 46.08) Hz; LM^sink^, 27.57 (24.65 – 30.57) Hz; p < 10^-5^, paired bootstrap test). **(H)** Mean difference in the interneuron to principal cell firing ratio between LM^sink^ and Rad^sink^ ripples. Note that while this ratio is similar between Rad^sink^ and LM^sink^ ripples in CA1 (p = 0.48; paired bootstrap test), CA3 exhibits a significantly higher ratio (i.e., higher interneuron firing relative to principal cells) in LM^sink^ ripples compared to Rad^sink^ ripples (p < 10^-5^; paired bootstrap test). **(I)** Firing response similarity of principal cells to Rad^sink^ versus LM^sink^ ripples for another example sleep session. Each point represents the firing rate of a CA1 (purple) or a CA3 (green) cell. The Pearson correlation coefficient (r) reports the similarity between the firing rates in Rad^sink^ versus LM^sink^ ripples, which we referred to as “response rate similarity.” Dashed purple and green lines represent the best-fit linear relationships for CA1 and CA3, respectively. **(J)** Similar to (I) but showing the spiking probability for the same two recording sessions as in Figure 2C and (I). **(K)** Mean difference in spiking probability similarity between CA1 and CA3 principal cells, expressed as a percentage relative to CA3. This indicates that the spiking probability similarity across Rad^sink^ and LM^sink^ ripples is significantly higher in CA1, suggesting that CA3 cells respond more differently between the two ripple types (p < 10^-5^; bootstrap test). **(L-N)** Classification accuracy of Rad^sink^ and LM^sink^ ripples in CA1 and CA3. Shown in (L) is a schematic of the method used to classify Rad^sink^ versus LM^sink^ ripples based on the entire spike train around ripples (see section ‘Population activity discrimination of Rad^sink^ versus LM^sink^ ripples’). The first two principal components (PCs) of these time responses were computed for each cell independently. PCs from all cells were then concatenated and used to train a logistic regression classifier to predict whether a given ripple was Rad^sink^ or LM^sink^. Shown in (M) is the classifier accuracy, as a function of the number of cells used for the training stage, separately for CA1 and CA3. Shown in (N) is the corresponding estimation plot representing the effect size for the difference in the accuracy of the CA1 versus CA3 classifiers (trained on 15 principal cells). CA3 discriminated Rad^sink^ versus LM^sink^ significantly better than CA1; p < 10^-5^; bootstrap test; n = 129 sleep sessions for CA1 and n = 89 for CA3; see also Table S2). This finding aligns with the observation that CA3 principal cells exhibit greater differences in response between Rad^sink^ and LM^sink^ ripples compared to CA1 (Figure 2). **(O-Q)** Ripple-type differences in coactivity and topological organisation. **(O)** Scatter plots showing the mean coactivity of CA1 (purple) and CA3 (green) principal cells during LM^sink^ versus Rad^sink^ ripples from two example sleep sessions. Each point represents the average coactivity value of a CA1 or a CA3 principal cell. The dashed black line indicates the diagonal of the first and third quadrants. Note that CA1 principal cells exhibit higher coactivity during LM^sink^ ripples, whereas CA3 principal cells show higher coactivity during Rad^sink^ ripples (all Ps < 0.01; bootstrap tests; sleep session 1: n = 21 CA1 and n = 17 CA3 principal cells; sleep session 2: n = 48 CA1 and n = 25 CA3 principal cells). **(P)** Mean difference of CA3 principal cells coactivity between Rad^sink^ and LM^sink^ ripples, expressed as a percentage change relative to Rad^sink^ ripples. The vertical red dashed line represents the mean coactivity in Rad^sink^ ripples. These results indicate, as highlighted by the black arrow, that CA3 principal cells are more coactive during Rad^sink^ ripples than LM^sink^ ripples, contrasting with the pattern observed for CA1 cells (Figure 3B; p = 2×10^-5^; paired bootstrap test). **(Q)** Mean difference in the structural balance of LM^sink^ coactivity graphs of CA3 principal cells relative to Rad^sink^ graphs. The vertical red dashed line marks the mean structural balance in Rad^sink^ ripples. There was no significant difference in structural balance between the two ripple types for CA3 coactivity graphs, unlike for CA1 graphs (Figure 3E; p = 0.53; paired bootstrap test). *p < 0.05, **p < 0.01, ***p < 0.001.

**Figure S5:**
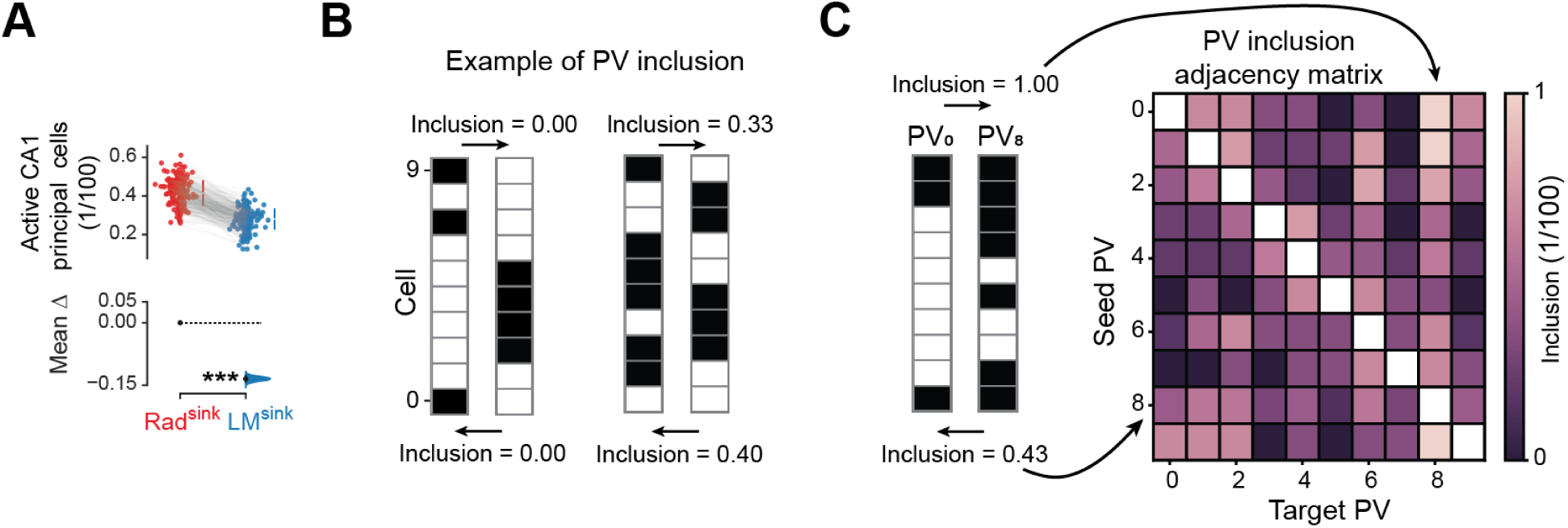
CA1 recruitment and inclusion analysis between Rad^sink^ and LM^sink^ ripples. **(A)** Estimation plot showing the effect size for the difference in the proportion of active CA1 principal cells (i.e., cells that fire at least one action potential) between the two ripple profiles. Rad^sink^ ripples recruit a higher proportion of CA1 principal cells compared to LM^sink^ ripples. Each dot represents a single sleep session. **(B,C)** Inclusion analysis. Shown in (B) is a schematic illustrating the procedure for calculating the inclusion of the set of active cells across two population vectors (PVs; one as the seed vector; the other as the target vector). Each active cell in each PV is depicted by a black field. Pairwise inclusion values are computed for each pair of PVs (*m*, *n*). Shown in (C) is an example adjacency matrix containing these inclusion values. *** p < 0.001.

**Figure S6:**
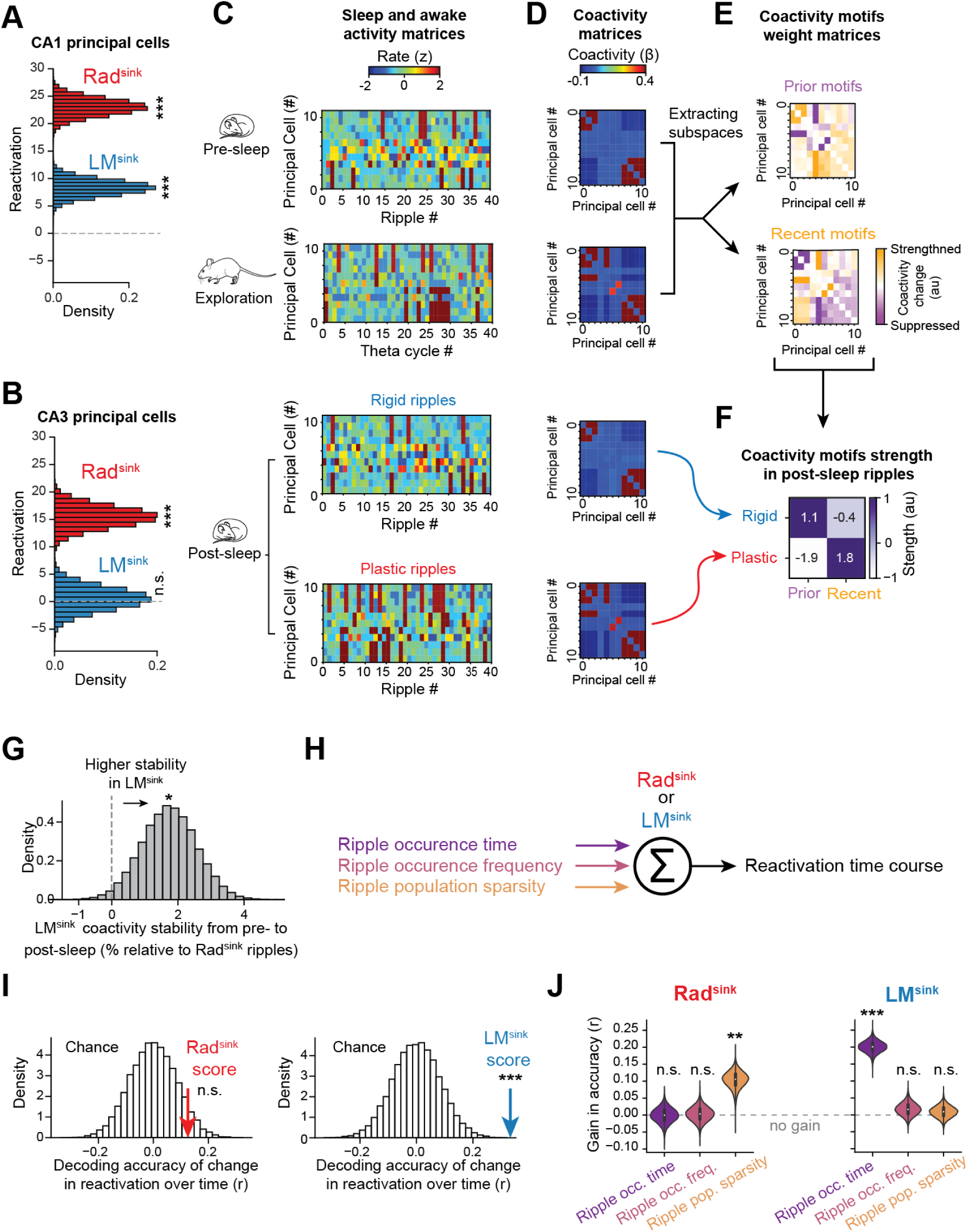
Reactivation of CA1 versus CA3 neurons and prior versus recent coactivity motifs. (A,B) Offline reactivation of waking theta patterns in CA1 and CA3 principal cells. Shown in (A) is the mean difference in the reactivation of CA1 principal cells during Rad^sink^ and LM^sink^ ripples, expressed as a percentage relative to pre-sleep (all Ps < 10^-5^; 1-tailed bootstrap tests). The same is shown in (B) but for CA3 principal cells, which reactivate in Rad^sink^ ripples (p < 10^-5^; 1-tailed bootstrap test) but not in LM^sink^ ripples (p = 0.39; 1-tailed bootstrap test) **(C-F)** Extraction and analysis of prior and recent coactivity motifs in simulated neural activity. We validated the method described in the section “Reactivation of prior and recent coactivity motifs” using an *in-silico* approach. We first simulated a pre-sleep population activity matrix containing two pre-defined motifs of coactive neurons (“prior motifs”). Subsequently (i.e., during awake exploration), one of these prior motifs underwent re-organisation by gaining an additional neuron, while a new (third) motif of coactive neurons emerged (“recent motif”). We then simulated two post-exploration sleep population activity matrices: one only expressed the motifs previously observed in pre-exploration sleep (“rigid ripples”; see section “Simulation of prior and recent coactivity motifs”), and the other incorporated the exploration-related motif (“plastic ripples”). Shown are example activity matrices over time bins for each session (C), with the corresponding coactivity matrices (D). We next extracted two subspaces from the pre-exploration sleep and the exploration coactivity matrices: the prior coactivity motifs (present in pre-sleep) and the recent motif (emerging during exploration and absent in pre-sleep) (E). These transformation matrices featured the cell pairs composing these motifs, highlighted in orange to denote strengthened connections (top panels for prior motifs and bottom panels for recent motifs). These transformation matrices were then applied to the simulated post-exploration sleep activity to evaluate the strength of each motif (F). The heatmap in (F) summarizes the strength of prior and recent coactivity motifs in the two post-exploration sleep cases from (C,D), demonstrating that rigid ripples more strongly reactivated prior motifs, while plastic ripples predominantly reactivated recently-expressed coactivity motifs. **(G)** Mean difference in the stability of CA1 principal cell coactivity from pre-exploration to post-exploration sleep in Rad^sink^ and LM^sink^ ripples (expressed as a percentage relative to Rad^sink^ ripples). LM^sink^ motifs are more stable than Rad^sink^, being more consistent across the two sleep sessions either side of exploration [mean stability in LM^sink^ ripples relative to Rad^sink^ ripples (95% CI): 1.74 (0.12 – 3.35) %; p = 0.018; 1-tailed paired bootstrap test]. **(H-J)** Factors explaining temporal changes in reactivation over hour-long sleep. We trained linear regression models to predict the reactivation strength of the recent motifs over the hour-long sleep/rest (H). These models were trained independently for Rad^sink^ and LM^sink^ ripples. Three features were used: ripple occurrence (occ.) time, ripple occurrence frequency (freq.), and ripple population sparsity as predictors (in 10-minute bins). Shown in (I) is the accuracy of each model alongside its corresponding chance level (*left*, Rad^sink^; *right*, LM^sink^). Chance level estimated by shuffling reactivation values 25,000 times. Red/Blue arrow indicates the observed model accuracy. Note that the Rad^sink^ model did not significantly predict the reactivation time course (p = 0.068; 1-tailed bootstrap test), consistent with the stable reactivation observed throughout sleep/rest (Figure 5F). In constrast, changes in reactivation over time during LM^sink^ ripples were significantly predicted (p < 4×10^-5^; 1-tailed bootstrap test). Shown in (J) are violin plots showing the contribution of each predictor to models’ accuracy. *Left*, for the Rad^sink^ model, only ripple population sparsity significantly contributed (p = 2.4×10^-3^, 1-tailed bootstrap test, Bonferroni corrected). *Right*, for the LM^sink^ model, only ripple occurrence time significantly contributed (p < 4×10^-5^, 1-tailed bootstrap test, Bonferroni corrected). *p < 0.05, **p < 0.01, ***p < 0.001.

**Supplementary Table S1.** Silicon probe details.

| Strain | RRID | n | Implant |
| --- | --- | --- | --- |
| C57Bl6/J | IMSR_JAX:000664 | 1 | NeuroNexus A1x32-5mm-25-177-H32_21mm |
| C57Bl6/J | IMSR_JAX:000664 | 1 | NeuroNexus A1x64-edge-6mm-20-177-H64LP_30mm |
| C57Bl6/J | IMSR_JAX:000664 | 3 | Cambridge NeuroTech ASSY-236 H3 Chronic 64-Molex |

**Supplementary Table S2.**
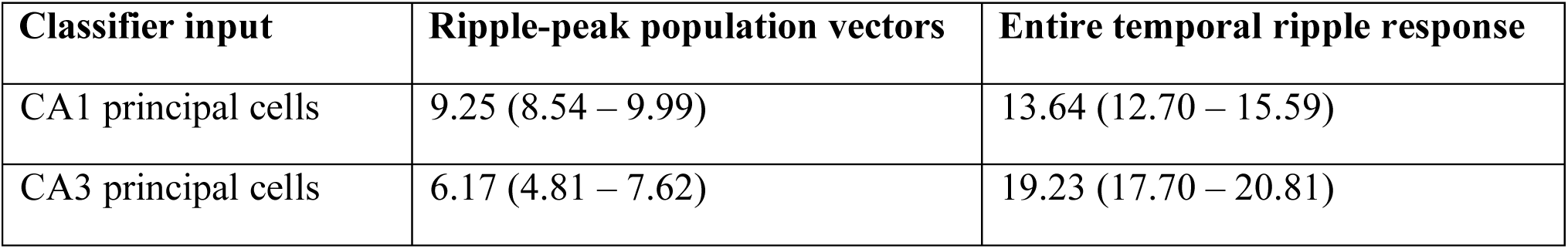
Performances of models discriminating Rad^sink^ versus LM^sink^. . Shown are the cross-validated performance [mean (95% CI); bits; x 10^-2^] of classifiers trained to predict Rad^sink^ versus LM^sink^ ripple types based on CA1 or CA3 activity, using either their population vectors (15 principal cells each) at the ripple peak or the entire temporal spiking profile around the ripple. Performance quantified as the mutual information (M.I.) between the true and predicted ripple types.

| Classifier input | Ripple-peak population vectors | Entire temporal ripple response |
| --- | --- | --- |
| CA1 principal cells | 9.25 (8.54 – 9.99) | 13.64 (12.70 – 15.59) |
| CA3 principal cells | 6.17 (4.81 – 7.62) | 19.23 (17.70 – 20.81) |

